# Integrative experimental/computational approach establishes active cellular protrusion as the primary driving force of phagocytic spreading by immune cells

**DOI:** 10.1101/2022.03.01.482589

**Authors:** Emmet A. Francis, Volkmar Heinrich

**Affiliations:** Department of Biomedical Engineering, University of California Davis, Davis, CA, USA

## Abstract

The dynamic interplay between cell adhesion and protrusion is a critical determinant of many forms of cell motility. When modeling cell spreading on adhesive surfaces, traditional mathematical treatments often consider passive cell adhesion as the primary, if not exclusive, mechanistic driving force of this cellular motion. To better assess the contribution of active cytoskeletal protrusion to immune-cell spreading during phagocytosis, we here develop a computational framework that allows us to optionally investigate both purely adhesive spreading (“Brownian zipper hypothesis”) as well as protrusion-dominated spreading (“protrusive zipper hypothesis”). We model the cell as an axisymmetric body of highly viscous fluid surrounded by a cortex with uniform surface tension and incorporate as potential driving forces of cell spreading an attractive stress due to receptor-ligand binding and an outward normal stress representing cytoskeletal protrusion, both acting on the cell boundary. We leverage various model predictions against the results of a directly related experimental companion study of human neutrophil phagocytic spreading on substrates coated with different densities of antibodies. We find that the concept of adhesion-driven spreading is incompatible with experimental results such as the independence of the cell-spreading speed on the density of immobilized antibodies. In contrast, the protrusive zipper model agrees well with experimental findings and, when adapted to simulate cell spreading on discrete adhesion sites, it also reproduces the observed positive correlation between antibody density and maximum cell-substrate contact area. Together, our integrative experimental/computational approach shows that phagocytic spreading is driven by cellular protrusion, and that the extent of spreading is limited by the density of adhesion sites.

**Author Summary:** To accomplish many routine biological tasks, cells must rapidly spread over different types of surfaces. Here, we examine the biophysical underpinnings of immune cell spreading during phagocytosis, the process by which white blood cells such as neutrophils engulf pathogens or other foreign objects. Our computational framework models the case in which a human neutrophil spreads over a flat surface coated with antibodies, which we also test experimentally in a companion paper. Our primary purpose is to assess whether phagocytic spreading is actively driven by protrusive forces exerted by the cell, or passively by adhesive forces acting between receptors in the cell membrane and antibodies on the surface. By directly comparing our model predictions to experimental results, we demonstrate that phagocytic spreading is primarily driven by protrusion, but the extent of spreading is still limited by the availability of binding sites. Our findings improve the fundamental understanding of phagocytosis and may also pave the way for future investigations of the balance between adhesion and protrusion in other forms of cell spreading, such as wound healing or cancer cell migration.

## Introduction

The ability of cells to spread on surfaces is critical to many biological processes, including cell migration, wound healing, and tissue formation. Immune cells are especially adept at rapid spreading; for instance, neutrophils commonly travel up to an order of magnitude faster in motility assays than fibroblasts or endothelial cells [1–5]. Vital immune cell functions, such as firm arrest at the endothelium and phagocytosis of pathogens, rely on this ability. A deeper understanding of the physical mechanisms driving cell spreading is essential for informed pharmaceutical strategies and new therapies targeting immune cells, such as current efforts to enhance phagocytosis for more effective clearance of cancer cells [6, 7].

Many experimental and theoretical studies have sought to unravel the physical principles of cell spreading by analyzing the initial growth of cell contact regions on a flat, adhesive substrate [8–11]. However, it often remains unclear whether any given type of cell spreading is primarily driven by passive forces due to adhesion or by active protrusive and/or contractile forces generated by the cell. In the simplest model, cell spreading is treated as a purely passive process, similar to a viscous droplet spreading on an adhesive substrate [8, 12, 13]. Alternative models assume that cytoskeleton-driven protrusion determines the rate of cell spreading, neglecting the effective attractive pre-contact force due to ligand-receptor interactions [10, 14, 15].

Here, we aim to illuminate fundamental cell-spreading mechanisms by focusing on the mechanistic driving force of phagocytic spreading. In conventional phagocytosis, an immune cell first adheres to the surface of a target particle such as a bacterium, often by binding to opsonic ligands such as immunoglobulin G (IgG) antibodies. The cell then spreads over the target surface until its membrane closes around the particle. On the other hand, when presented with an excessively large target, the cell is unable to fully engulf the particle and continues to spread until its target-contact area reaches a maximum [16, 17]. Such cases, including phagocytic spreading on flat substrates, are termed frustrated phagocytosis. The relatively simple geometry of a cell undergoing frustrated phagocytic spreading on a flat surface is particularly amenable to theoretical analysis, especially when the cell spreads in an axisymmetric manner. Moreover, it seems reasonable to assume that some of the insights generated by integrative experimental/theoretical analyses of cell spreading on flat surfaces also carry over to other geometries [18–21]. For example, a previous mathematical model of phagocytosis has been able to predict experimentally observed variations in the engulfment of spherical particles in the size range of 3-11μm by changing only one model parameter value, i.e., the input size of the target bead [22].

Reflecting the above-mentioned contrasting mechanistic notions about the driving force of generic cell spreading, two hypotheses have been put forward to explain phagocytic spreading, i.e., the “Brownian zipper” and “protrusive zipper” models [23]. Both models concur that fresh contact between the cell membrane and target surface results in a zipper-like adhesive attachment that is essentially irreversible, in agreement with experimental observations [16, 24, 25]. However, regarding the driving force of cellular advancement, the Brownian zipper model postulates that strong adhesion alone is responsible for pulling the cell membrane onto the target surface. In contrast, the protrusive zipper model assumes that an active protrusive force generated by the cytoskeleton is the predominant driver of outward motion of the cell surface, without excluding a possible contribution of adhesive cell-substrate interactions. It is important to bear in mind that our definition of these two models in terms of the driving force of spreading is not the same as the distinction between “passive” and “active zippers” used elsewhere [26]. In contrast, neither of the zipper models defined in [26] included an active protrusive force in the computations but instead implemented an effective adhesion stress as the sole driving force of spreading. Therefore, both zipper models defined in [26] fall into the category of a Brownian zipper as defined here.

We seek to establish which of these hypotheses is more realistic by developing mathematical models of the mechanics of isotropic cell spreading and comparing our theoretical predictions with experimental observations of human neutrophils spreading on flat, IgG-coated surfaces. As reported in detail in a companion paper, our experimental design allowed us to examine the spreading behavior of neutrophils on surfaces coated with different densities of IgG [24]. Because higher IgG densities correspond to stronger adhesive forces, we reasoned that the Brownian zipper model should predict a much stronger correlation between IgG density and rate of contact-area growth (“spreading speed”) than the protrusive zipper model. Here, we use computer simulations of phagocytic spreading to verify this common-sense expectation and further investigate mechanisms governing immune-cell spreading. First, we present the predictions of a Brownian zipper model that realistically implements known insights about the mechanical behavior of human neutrophils. Then, we investigate the protrusive zipper model, including the contribution of adhesive pre-contact interactions to the driving force of spreading. Finally, we also present a version of the Brownian zipper model that implements the discrete nature of receptor-ligand interactions more realistically than the continuum models introduced earlier. Together, our integrative experimental/theoretical approach establishes that the Brownian zipper model is inconsistent with experimental results, whereas the spreading dynamics predicted by the protrusive zipper model agree well with our experiments.

## Results

### Model implementation of key biophysical mechanisms of cell spreading

We consider a simple model cell that captures the basic geometrical and mechanical properties of human neutrophils, the most abundant type of white blood cell and one of the prototypes of professional phagocytes. This model cell consists of an axisymmetric, highly viscous fluid body of constant volume that is surrounded by an elastic cortex with uniform tension (Fig 1). Termed the “cortical shell, liquid core model”, this theoretical concept has successfully reproduced the passive response of human neutrophils to imposed deformations [27, 28]. In the undeformed reference state, the persistent tension of the cortex maintains a spherical cell shape with a typical diameter of 8.5 μm. The nucleus of neutrophils is segmented into two to five lobes [29, 30] and appears to play only a minor role during deformations, lending support to the approximation of the cell interior as a homogeneous fluid. The effective viscosity (*μ*) of the cytoplasm has been found to be about 200,000 times the viscosity of water at room temperature in passive neutrophils [27, 28], and another ~3-times higher in active neutrophils [31]. It is well known to increase during phagocytosis [22, 32, 33]. We use a viscosity value of *μ* = 200 Pa-s in our passive Brownian zipper model and a value of *μ* = 1660 Pa-s in our active protrusive zipper model. The original assumption of a constant cortical tension [27] was later found to be approximately true only for small deformations [34]. The revised relationship between tension (*τ*) and cell surface area (*A_cell_*) was shown to be roughly biphasic [35]. Assuming an isotropic and uniform tension, our simulations incorporate a smoothed version of the following biphasic constitutive relationship, in overall agreement with a number of quantitative experimental studies [22, 35, 36] (plotted in S6 Appendices, Appendix C):

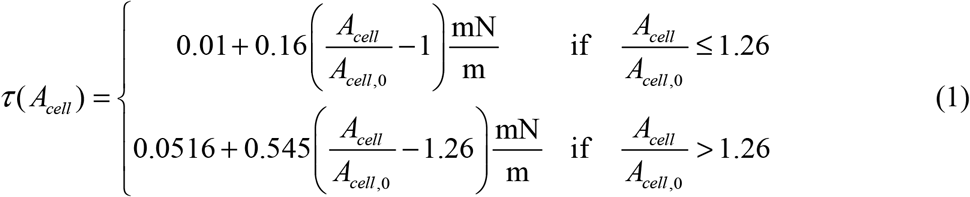

where *A_cell,0_* is the surface area of the resting neutrophil before spreading. We also tested alternative versions of this constitutive relationship in S6 Appendices, Appendix C, and found that they do not qualitatively alter the results of our simulations.

**Fig 1.**
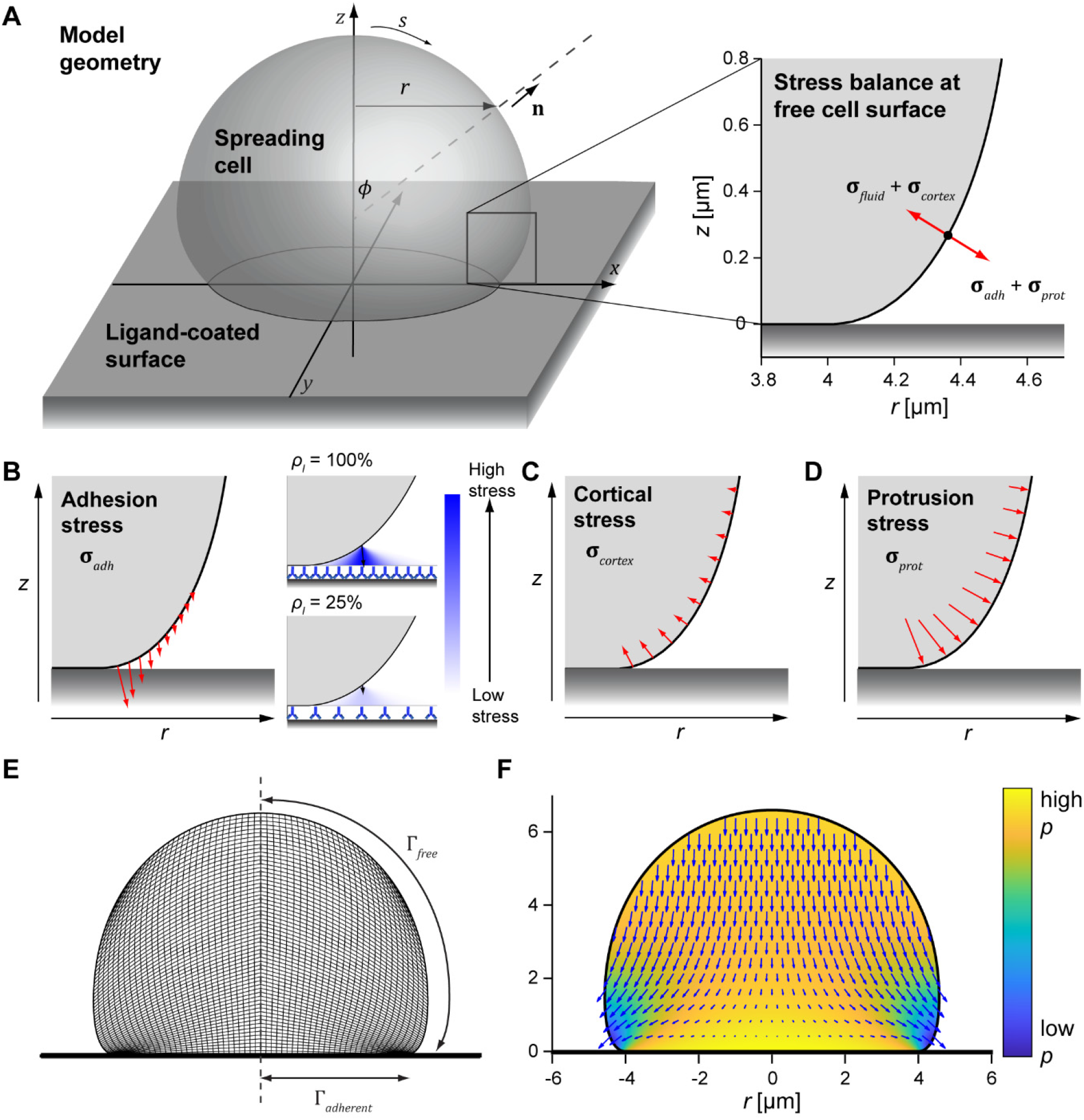
Illustration of our computational model of isotropic cell spreading. (A) 3D rendering shows the geometry and defines coordinates for an axisymmetric spreading cell on a flat surface. The enlarged inset (right) illustrates the stress balance at the free cell boundary. (B) Pre-contact adhesion stress effectively pulls the membrane onto the flat surface. The enlarged insets conceptually depict how the adhesion stress is computed for different ligand densities *ρ_l_*. (C) Cortical tension and membrane curvature give rise to an effective inward normal stress. (D) The protrusion stress acts normal to the cell membrane and is concentrated at the region of the membrane closest to the substrate. (E) The cross-sectional snapshot of a simulation illustrates the mesh composed of quadrilateral elements used in the calculations, with tighter element packing closer to the flat surface. The computational domain only includes half of this mesh due to axial symmetry. Boundary regions are labeled Γ, and the vertical dashed line is the symmetry axis (*r* = 0) (F) This snapshot of a simulation illustrates the fluid velocity (vector field) and relative pressure (heat map) computed for a given cell shape with known boundary stresses.

Movements of the viscous cytoplasm generally do not exceed velocities of 1 μm/s during phagocytic cell spreading and are characterized by low Reynolds numbers. Such movements are well described by the Stokes equations for creeping flow:

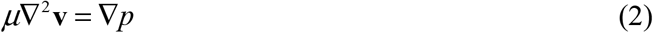

along with the continuity equation for incompressible fluids:

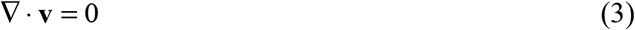

Here, **v** denotes the fluid velocity vector, and *p* is fluid pressure. Together, these equations govern the flow behavior of our model cell.

Interactions between the cell surface and the substrate can facilitate cell spreading directly via adhesion, and indirectly through receptor-induced signaling that leads to protrusion. Including the respective stresses, the moving boundary condition governing the free (i.e., nonadherent) part of the cell surface (Γ*_free_*) balances up to four types of stress vectors: fluid stress (**σ***_fluid_*), adhesion stress (**σ***_adh_*), protrusion stress (**σ***_prot_*), and cortical stress (**σ***_cortex_*) (Fig 1A):

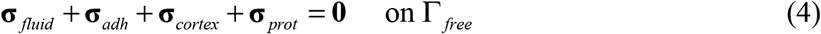

The fluid stress vector **σ***_fluid_* encompasses the pressure difference across the cell surface as well as the stress exerted on the surface by the moving cytoplasm. It is determined by the constitutive relation for a Newtonian fluid:

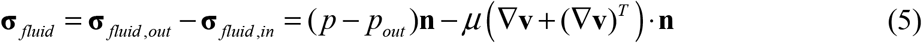

where **n** is the unit surface normal, and the dot in the last term on the right-hand side denotes the inner product between a tensor and a vector. The extracellular medium has a much lower viscosity than the intracellular fluid; therefore, Eq (5) neglects the medium’s viscous contribution and only accounts for its hydrostatic pressure denoted *p_out_*.

Our implementation of the adhesion stress is based on the assumption that adhesion can effectively be represented by a short-range attractive potential between the cell surface and the ligand-coated substrate. We compute the effective adhesion stress at a given point on the cell surface by summing over all pairwise interactions with substrate molecules that lie outside the cell-substrate contact region at the time (Fig 1B). For a continuum of ligand molecules, the resulting expression takes the form (derived in S6 Appendices, Appendix A)

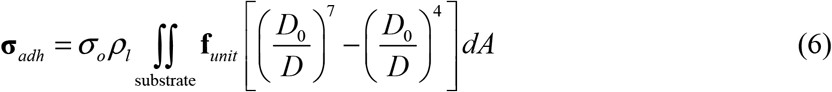

where *ρ_l_* is the ligand surface density and *σ_0_* is a scaling factor representing the strength of interaction per unit area of the cell surface. The unit vector **f***_unit_* specifies the direction of a given interaction, *D* denotes the distance between the interacting points, and *D_0_* is a constant setting the overall range of the adhesion potential (set to 50 nm in our simulations; S7 Table).

The origin of the cortical stress is the cortical tension, which resists expansion of the cell surface area and acts along the curved surface of the cell (Fig 1D). The resulting stress acts normal to the surface and is given by the product of the tension *τ* (Eq (1)) and the mean curvature of the cell surface, as dictated by Laplace’s law:

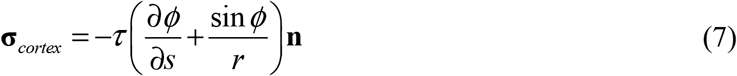

Here, the term in parentheses is the mean curvature of the axisymmetric cell surface expressed in the coordinates defined in Fig 1A.

The protrusion stress comprises outward forces exerted on the cell surface by the actively remodeling cytoskeleton. Purposeful reorganization of intracellular structures requires tight coordination by biochemical signaling cues, which in turn are induced by receptor-ligand binding. Rather than attempting to implement these highly complex mechano-chemical mechanisms, we capture their localized character in a semiempirical manner. Our model postulates that the protrusive stress is greatest at the perimeter of the cell-substrate contact region and decays as a function of the distance from this leading edge measured along the arc length (*s*, Fig 1A) of the contour of the unbound cell surface. We assume that this stress acts normal to the cell boundary, and that its decay along the free cell surface is exponential (Fig 1C):

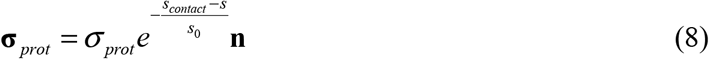

In this equation, *s_contact_* is the arc length value at which the cell-surface contour makes contact with the substrate, and *s_0_* is the characteristic length that sets the spatial range of the protrusion stress.

Bearing in mind that cytoskeletal remodeling also is responsible for the behavior of the cortical tension, and that the tension varies during cell spreading (Eq (1)), it is reasonable to assume that the overall strength of the protrusion stress (*σ_prot_*) varies in a similar fashion. In addition to the spatial distribution of the cortical stress governed by Eq (8), we therefore also postulate a deformation-dependent strength of the protrusive stress. Our choice of this dependence, and its motivation, will be explained in a later section.

Finally, for the part of the cell surface that is in contact with the substrate, we enforce a no-slip boundary condition, requiring that

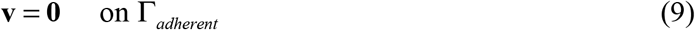

Each simulation begins with a model cell that rests on the surface and has formed a very small initial contact footprint (area < 0.1 μm^2^). The shape of the upper, free part of the cell at this time is a spherical cap. We determine the fluid velocities and pressure distribution at each time step by solving a perturbed form of the Stokes equations using the finite element method (FEM; Fig 1E,F) as described previously [37]. We then use the computed velocities to evolve the cell shape over a small time step. Details of our model implementation and solution method are given in Methods and S6 Appendices, Appendix D.

This generic modeling framework of cell spreading can easily be adopted to represent different mechanistic notions by adjusting the relative contributions of the adhesion and protrusion stresses. Disregarding the protrusion stress altogether results in a mathematical representation of the Brownian zipper model described in the Introduction. On the other hand, the protrusive zipper model generally includes both the protrusion stress as well as the adhesion stress. A purely protrusion-driven mode of spreading also can be tested by setting the pre-contact adhesion stress to zero, although one needs to bear in mind that irreversible adhesion still maintains the cell-substrate contact region in this case. In all model versions, the adhesion stress can be varied over several orders of magnitude to predict how the spreading behavior depends on the density of ligand displayed on the substrate.

### Quantitative experimental results for the validation of model versions

The computational framework presented in the previous section can be adopted to simulate interactions by various cell types with targets of different axisymmetric geometries. Our present focus on human neutrophils spreading on flat substrates is motivated by a wealth of experimental results obtained for this particular scenario and presented in a companion paper [24]. In brief, we exposed cells to glass coverslips coated with IgG antibodies at densities (*ρ_IgG_*) spanning ~3 orders of magnitude. The analysis of these highly controlled experiments included measurements of the cell-substrate contact area of spreading cells over time (Fig 2). Regardless of IgG density, we found that the area-versus-time curves typically were sigmoidal in shape (Fig 2B,C). Defining a cell’s “spreading speed” as the fastest rate of contact-area growth (i.e., the slope of a sigmoidal fit to the area-versus-time curve at the inflection point), we found that all spreading cells increased their substrate-contact area at speeds of around 3 μm^2^/s irrespective of the density of deposited IgG (Fig. 2D). The measured values of the maximum contact area of spreading cells, on the other hand, exhibited a moderate but significant increase at higher IgG densities (Fig 2E). These experimental results provide useful quantitative benchmarks for comparison with computer simulations and will be used in the following sections to evaluate the validity of different mechanistic notions about cell spreading.

**Fig 2.**
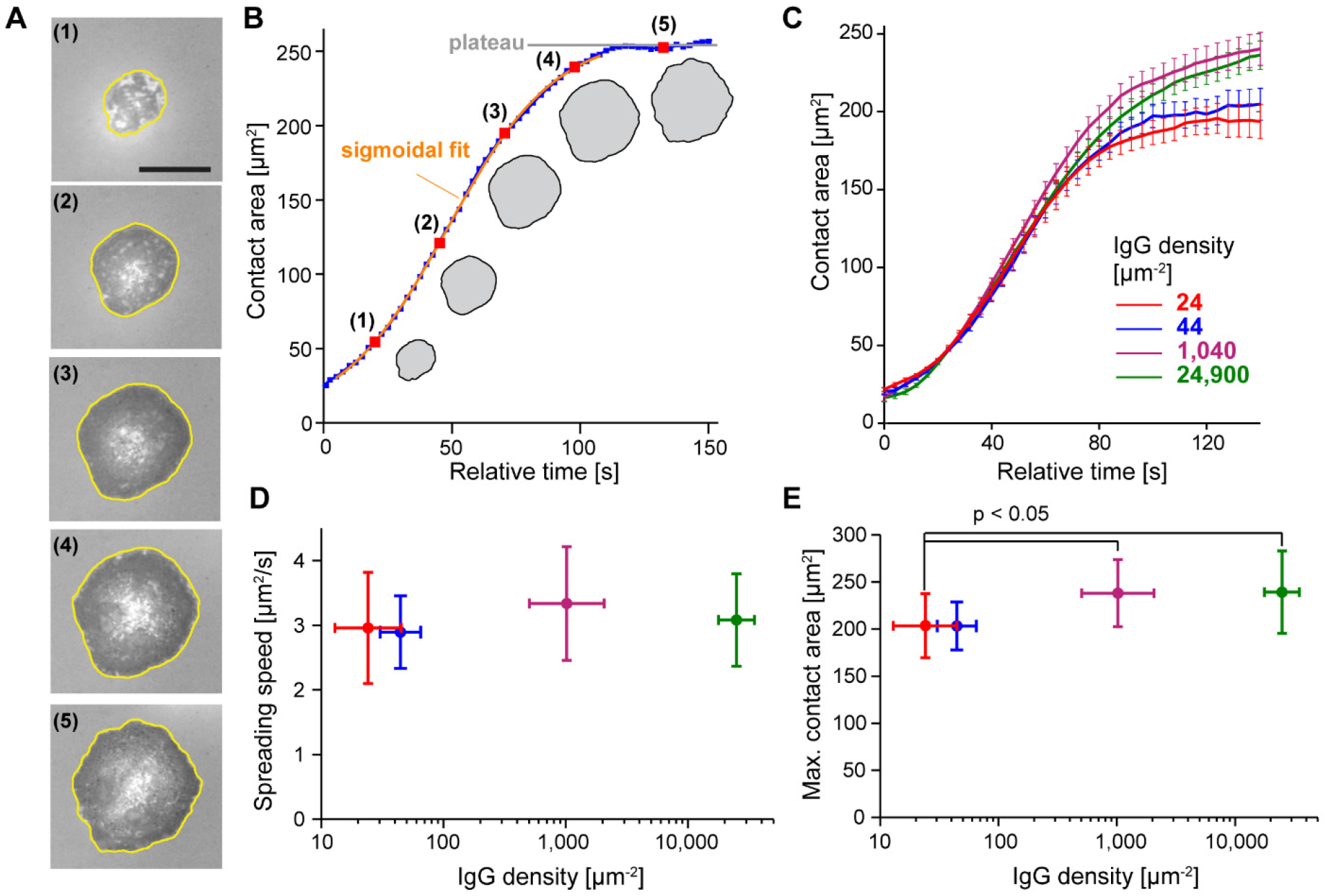
Summary of experimental results. (A) The contact region of a spreading neutrophil was imaged using reflection interference contrast microscopy (RICM). Contact regions in this example are traced in yellow. The scale bar in the first image denotes 10 μm. (B) Contact area is plotted as a function of time for the spreading cell shown in (A). The spreading speed is defined as the maximum slope extracted from a sigmoidal fit, and the maximum contact area is given by averaging the contact area at the plateau. (C) Mean contact area is computed at discrete time values for aligned curves of cells spreading on different densities of IgG. Mean quantified IgG densities are reported in the figure legend as IgG molecules per μm^2^. Error bars represent standard error of the mean. (D) Average spreading speed was quantified for the different IgG densities. There is no significant difference between mean slope values. (E) Maximum contact area increases slightly as a function of IgG density. Statistically significant differences in maximum contact area values were verified using ANOVA followed by a Tukey post hoc test. Error bars in both (D) and (E) indicate standard deviation. The IgG density was quantified for each condition as described in [24].

An additional measure of interest can be inferred from the actual surface densities of IgG used in the experiments. The most densely coated coverslips presented about 25,000 IgG molecules per μm^2^ and induced ~85% of deposited cells to spread. Remarkably, even at a low IgG surface density of ~44 molecules per μm^2^, about 30% of the deposited cells still spread in an IgG-dependent manner. Fcγ receptors bind IgG with an average equilibrium constant of about 10^−6^ M [38], corresponding to a binding energy per bond of *E_bind_* = −*k_B_T* ln (10^−6^) ≈ 14*k_B_T*. Assuming that all IgG molecules in the cell-substrate contact region have formed bonds with the cell’s Fcγ receptors, conservative (upper-limit) estimates of the adhesion energy density of cell-substrate interactions on substrates coated with the high and low numbers of IgG yield 1,500 μJ/m^2^ and 2.6 μJ/m^2^, respectively.

### Purely adhesion-driven spreading: predictions by the Brownian zipper model

We first tested the simplest version of our framework, the Brownian zipper model, by simulating purely adhesion-driven spreading in the absence of any protrusion stress (Fig 3, S1 Movie). In this scenario, the model cell is passive and is pulled onto the substrate by short-range attraction. This case is conceptually equivalent to a viscous droplet spreading on an adhesive substrate.

**Fig 3.**
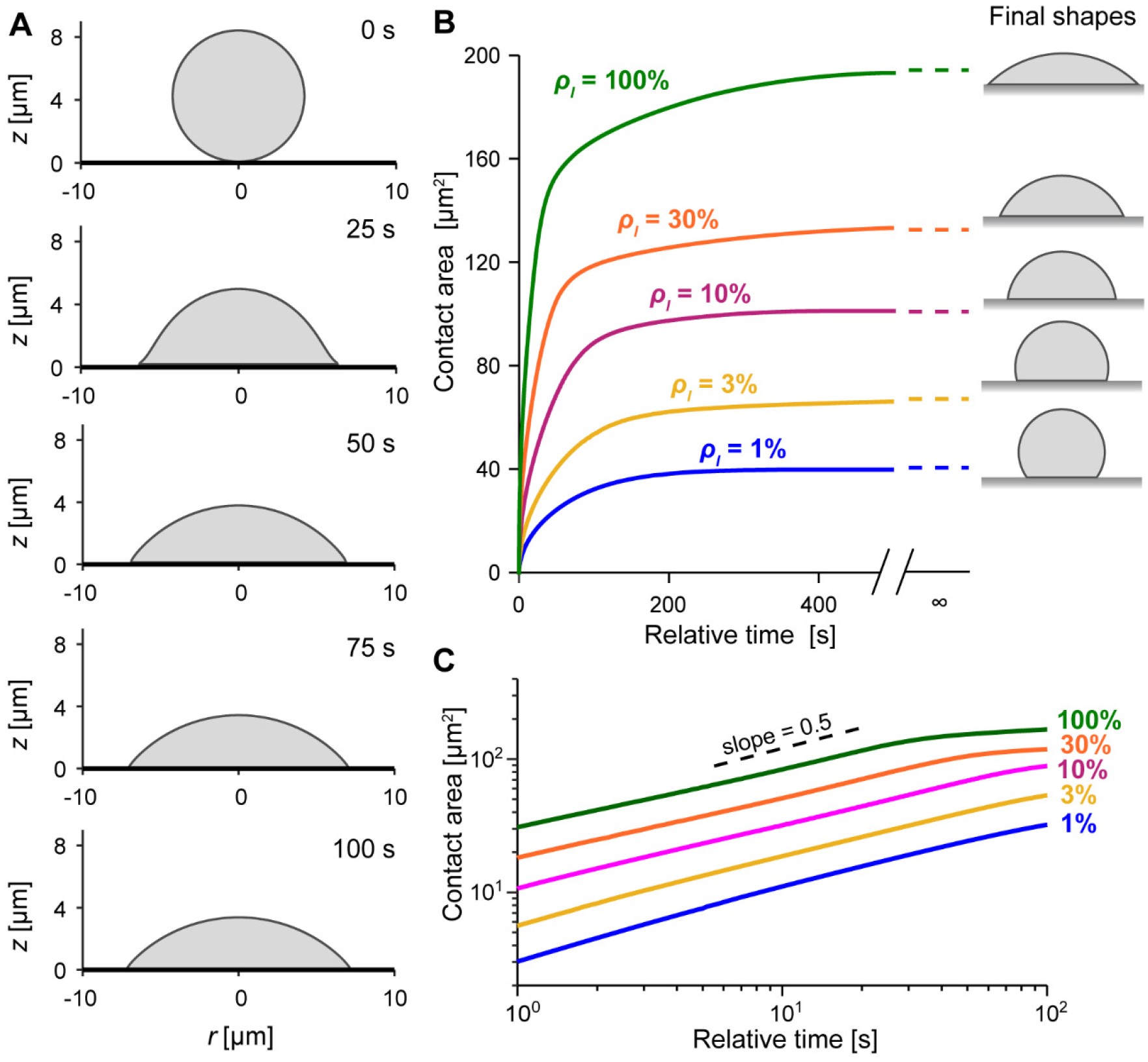
Predictions of the Brownian zipper model. (A) A model cell spreading on the highest tested ligand density (100%) quickly approaches a spherical cap morphology. Time stamps are included for each snapshot. (B) Curves of contact area vs. time show that the spreading rate decreases monotonically. The contact area ultimately approaches steady-state values predicted by the Young-Dupre equation (dashed lines). Predicted equilibrium shapes are included on the right. (C) Log-log plot of contact area vs. time is nearly linear over the initial spreading phase. The slope of this line corresponds to the exponent of a power law describing contact area growth as a function of time. The dashed line shows a slope of 0.5 for reference.

Varying the ligand density *ρ_l_* over two orders of magnitude, we found that the model cells generally spread fastest immediately after first contacting the substrate. Afterwards, the growth rate of the cell-substrate contact area decreased monotonically until the contact area reached a steady-state plateau marking equilibrium (Fig 3B). As expected, the equilibrium shapes were spherical caps. Log-log plots of the contact-area-versus-time predictions during the initial, fast spreading phase were approximately linear, with slopes close to 0.5 (Fig 3C). Hence, the contact-area growth initially obeys a power law, in agreement with various laws proposed to describe droplet wetting dynamics [39] or passive cell spreading [8, 13]. In fact, the exponent of 0.5 is an excellent match to results of a previous study that modeled cells as highly viscous droplets [13]. In S6 Appendices, Appendix B, we show how this exponent naturally arises during early spreading of a passive model cell driven by adhesion stresses concentrated at the contact perimeter.

Although the predictions of the Brownian zipper model agree well with the typical theoretical behavior expected for this kind of passive model cell, they clearly differ from the results of our phagocytic spreading experiments (Fig 2) in several important ways. First, the simulated contact-area-versus-time curves lack the characteristic sigmoidal shapes observed in experiments. Second, the predicted spreading curves exhibit a very strong dependence on the density of encountered ligands, in stark contrast to the observed spreading behavior. This dependence is evident in both the predicted spreading speeds, which increase dramatically with ligand density, as well as the predicted maximum contact areas, which increase several-fold as a function of the tested adhesive strengths.

Finally, the equilibrium shapes obtained for the adhesion energy densities used in the simulations are inconsistent with the cell deformations observed on the experimentally tested IgG densities. At equilibrium, the adhesion energy density (*γ*) of the Brownian zipper model cell is related to the cortical tension *τ* and the contact angle *θ_c_* via the Young-Dupre equation:

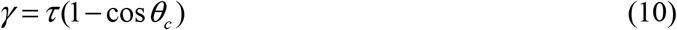

For a given value of the maximum cell-substrate contact area *A_c_*, the equilibrium shape of the model cell—a spherical cap—is uniquely defined by *A_c_* and the known, constant cell volume. Thus, basic geometry allows us to calculate the contact angle *θ_c_* as well as the total cell surface area *A_cell_*. Inserting the value of the latter into Eq (1) provides the cortical tension at equilibrium, which then can be used in Eq (10) to obtain the adhesion energy density as a function of *A_c_* (Fig 4). As expected, the relationship between maximum contact area and adhesion strength predicted by our numerical simulations agrees well with the Young-Dupre equation (S6 Appendices, Appendix A). The quantitative comparison of this theoretical prediction to actual adhesion energy densities—estimated from the densities of deposited IgG molecules as explained earlier—exposes entirely different behaviors between the model and experiments, however (Fig 4). Especially in experiments performed at lower IgG densities, the conservatively estimated adhesion energy densities were much lower than what would be required to form the measured maximum contact areas solely by adhesion. In summary, the results of this section rule out adhesion as the primary driving force of frustrated phagocytic spreading.

**Fig 4.**
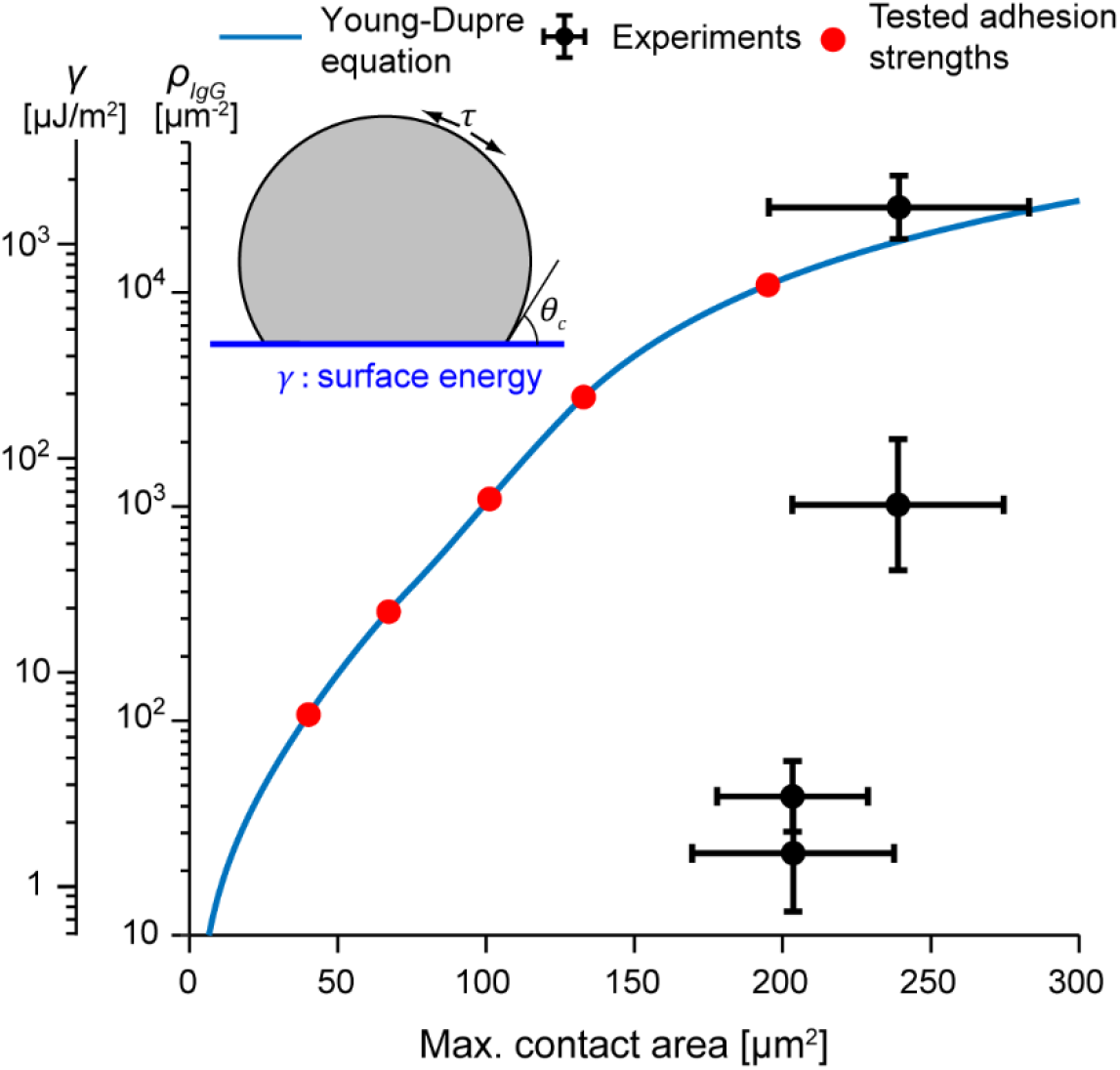
Relationship between ligand density and maximum contact area, as predicted by the Young-Dupre equation. The adhesion energy *γ*, and therefore the ligand density *ρ_IgG_*, can be expressed as a function of the contact area of the final spherical cap formed by a viscous droplet. The tested adhesion strengths are marked as red dots. Actual experimental values are shown for comparison. Error bars indicate standard deviation.

### Protrusion-dominated spreading: predictions of the protrusive zipper model

The protrusive zipper model incorporates the protrusion stress into the basic framework considered in the previous section, enabling us to account for active force generation by the model cell. Our semiempirical implementation is motivated by biological plausibility, and by the goal to reproduce experimental observations with a minimum set of rules for the spatial distribution (Eq (8)) and the deformation-dependent evolution of the protrusion stress during cell spreading. To capture the presumed correlation between receptor-induced signaling and protrusive force generation by the cytoskeleton, we express the strength of the protrusion stress as a function of the cell-substrate contact area *A_c_*. We assume that in the initial low-tension regime, the protrusion stress grows linearly with *A_c_*. Once the cortical tension increases more steeply, the protrusion stress is assumed to grow faster as well. Ultimately, the strength of the protrusion stress levels out at a maximum value of *σ_prot,max_* = 3.5 kPa, within the range of experimentally measured stresses exerted by parallel actin filaments [40, 41]. This behavior is implemented in our model through a smoothed version of the following rule (plotted in S6 Appendices, Appendix C):

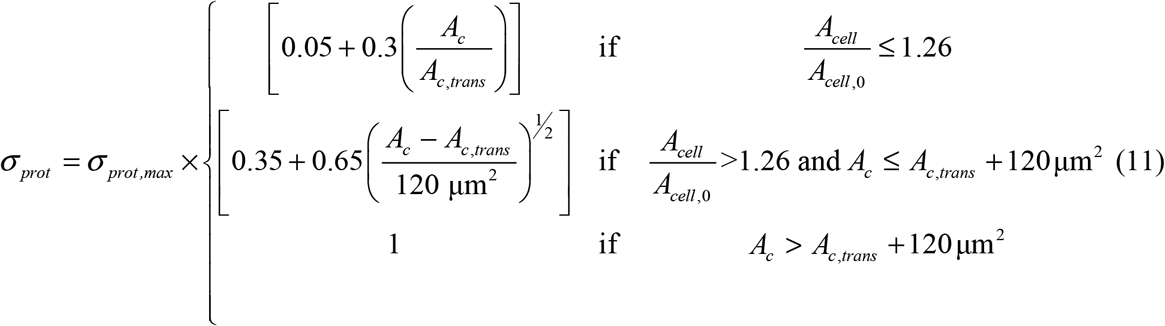

where *A_c,trans_* is the contact area when the total cell surface area *A_cell_* is equal to 1.26*A_cell,0_*, the transitional moment when the cortical tension starts rising more rapidly (Eq (1)).

We started examining the behavior of this model by first disregarding the adhesion stress altogether. In this limiting scenario, cell spreading is driven by protrusion, whereas the only remaining role of adhesion is to render cell-substrate contacts irreversible, as in all our models. As mentioned above, cell-substrate contact is coupled to protrusion through biochemical signaling via Eq 11. We found that in this case, the model cell begins spreading slowly while a distinctive leading edge gradually develops (Fig 5A, S2 Movie), resulting in a morphology that clearly differs from the spherical caps predicted by the Brownian zipper model (Fig 3). Spreading slows noticeably at about 90 s, around the time the protrusion stress reaches its maximum value. The resulting contact-area-versus-time curve has a sigmoidal shape, in agreement with the mean spreading behavior observed in experiments on the highest density of IgG (Fig 5B). We also tested slightly different versions of Eq 11 and found that the overall curve remains sigmoidal even for these alternative relationships (S6 Appendices, Appendix C). On average, spreading appears to commence somewhat more gradually in experiments than in our model, which may indicate an overestimate of the initial protrusion stress in Eq 11. Plots of the protrusion stress and the cortical tension as a function of time (Fig 5C) illustrate the qualitatively similar evolution of these two quantities, with the rise of the tension lagging due to the initial “slack” of the cell surface in the low-tension regime (Eq (1)).

**Fig 5.**
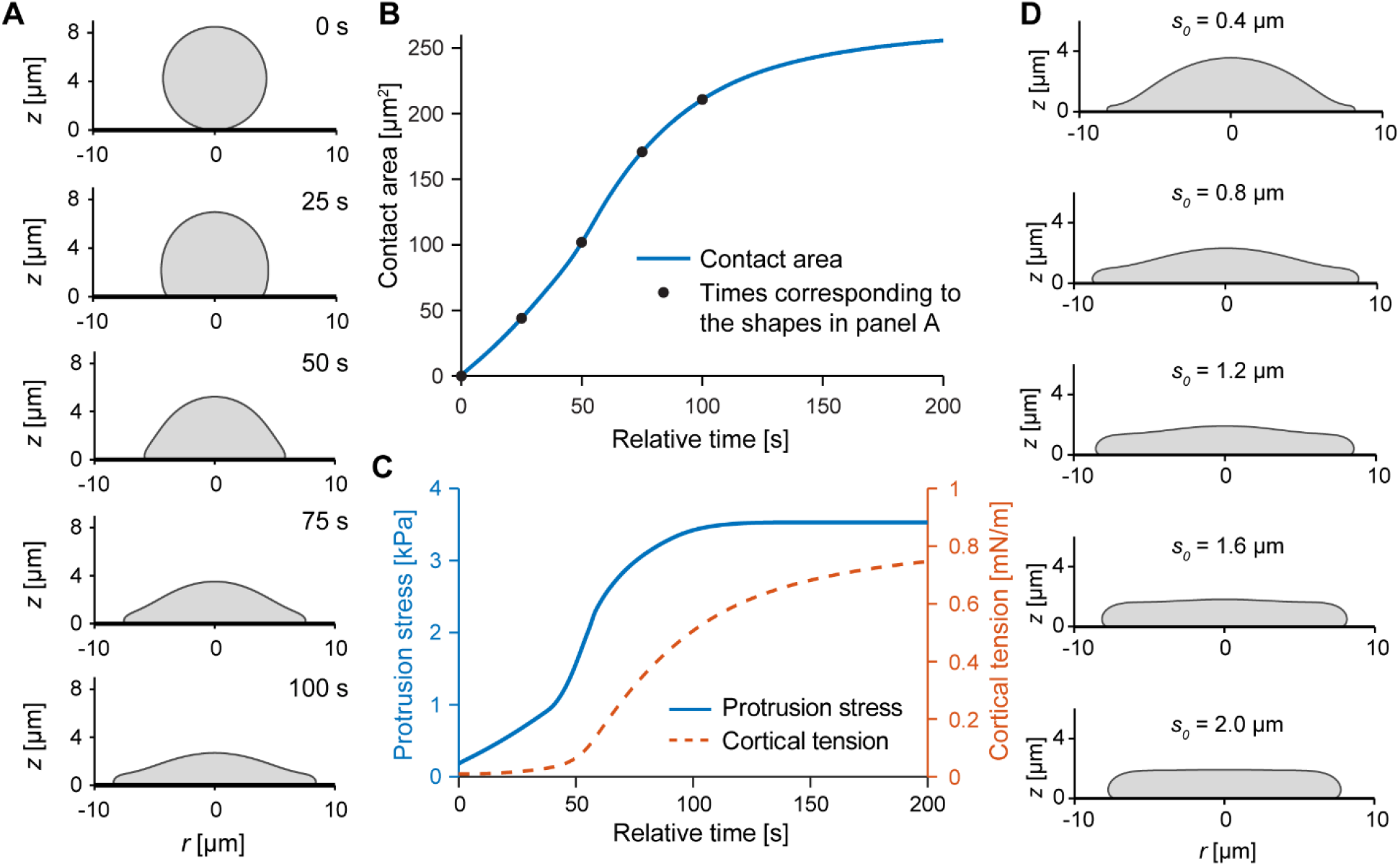
Predictions of the protrusive zipper model in the absence of adhesion stress. (A) For purely protrusion-driven spreading, lamellar pseudopods form as the protrusion stress increases over time. Time stamps of the the shown sample shapes are included. (B) Contact area increases sigmoidally over time for the protrusive zipper model, in good agreement with our measurements of the time course of the mean contact area on the highest density of IgG. Filled circles indicate where the shapes shown in panel A were computed. (C) Protrusion stress (governed by Eq (11)) and cortical tension (governed by Eq (1)) increase during cell spreading. (D) Varying the parameter *s_0_* determines the thickness of the leading lamella, resulting in a thin pseudopod for *s_0_* = 0.4 μm (top) and coordinated global cell deformation for *s_0_* = 2.0 μm (bottom). The sample shapes are shown at t = 120 s. All other protrusive zipper simulations in this paper were carried out using *s_0_* = 0.8 μm.

We chose the value *s_0_* = 0.8 μm for the characteristic length that defines the spatial range of the protrusion stress (Eq (8)) in the simulations shown in Fig 5A-C. Because *s_0_* does not appear to be a directly measurable quantity, it is instructive to inspect how its value affects the spreading model cell. Snapshots from simulations using different values of *s_0_* demonstrate how the cell morphology (here: at 120 s) depends on the spatial range over which the protrusion stress acts (Fig 5D, S3 Movie). The leading edge of the cell assumes the shape of a thin lamella for small values of *s_0_* and becomes increasingly thicker at larger *s_0_*-values. Aside from this varying thickness of cellular protrusions, the overall spreading behavior is similar for different values of *s_0_*.

Next, we inspected the behavior of the full protrusive zipper model by repeating the simulation with the same settings as used for Fig 5 but now also including the adhesion stress that captures the contribution of pre-contact attraction between cell and substrate to cell spreading. We again examined the same set of adhesion strengths as used in our tests of the Brownian zipper model (Fig 3). The simulations expose how, in this scenario, cell-substrate adhesion promotes spreading, affecting the overall spreading behavior in a quantitative manner but without altering it qualitatively. The model cells spread somewhat faster and farther on substrates coated with higher ligand densities (Fig 6A), with slightly larger contact angles between their leading edge and the substrate (Fig 6B, S4 Movie), but otherwise exhibited unchanged morphology. The sigmoidal character of the contact-area-versus-time curves is conserved through most of the tested adhesion strengths, and both the spreading speeds as well as the maximum contact areas vary much less than in the predictions of the Brownian zipper model (Fig 3).

**Fig 6.**
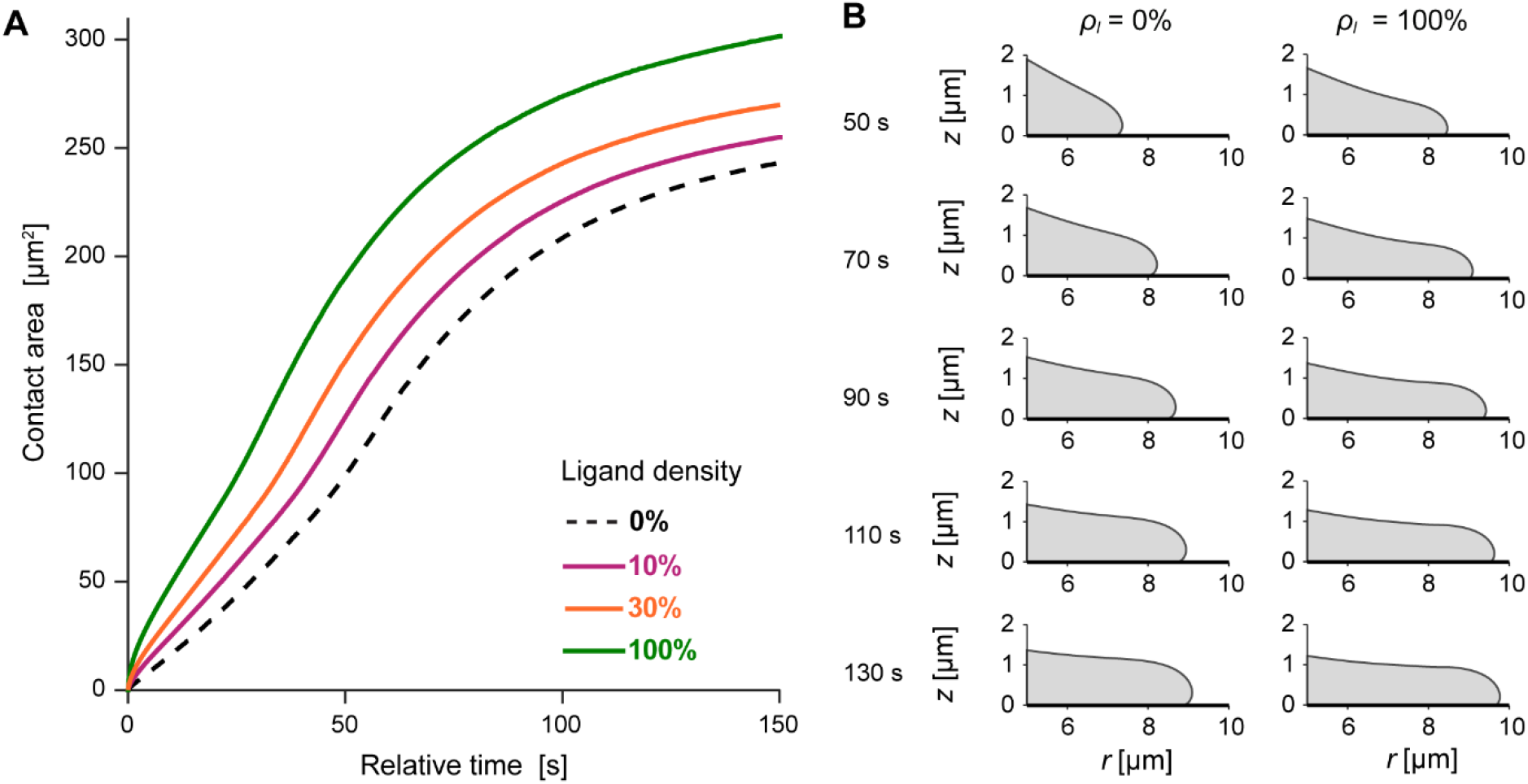
Predictions of the protrusive zipper model with varying adhesion stress. (A) Contact area increases faster and reaches higher values for higher ligand densities. The protrusion-only curve is shown for reference as a dashed line. (B) Magnified views of the leading edge of the model cell highlight subtle, adhesion-dependent changes of the pseudopod morphology. The shapes of the leading cell edge are shown for *ρ_l_* = 0% and *ρ_l_* = 100% at times ranging from 50 s to 130 s. On higher ligand density, both the extent of spreading and the dynamic contact angle increase. Other than that, the cell shapes exhibit little dependence on adhesion strength within the full protrusive zipper model.

Clearly, the predictions of the protrusive zipper model are in much better agreement with experimental observations than those of the Brownian zipper. A spreading cell morphology that is characterized by a forward-bulging leading lamella (Figs. 5–6) matches the cross-sectional profile of phagocytes engulfing large particles well [22, 25], in contrast to model cell shapes whose leading edge coincides with the perimeter of contact between the cell and substrate (Fig 3). The timing of spreading and sigmoidal shapes of the contact-area-versus-time curves predicted by the protrusive zipper model (Fig 5–6) agree with experiments as well, and the time-dependent increase in protrusion stress is consistent with a biphasic signaling response measured during FcγR-mediated phagocytosis [42]. Furthermore, both the predicted spreading speeds as well as the maximum contact areas are comparable to the respective values measured in experiments, and the slight increase of the maximum contact area at higher adhesion strengths is consistent with our measurements (Fig 6A). On the other hand, the predicted increase of the spreading speeds on higher ligand densities (Fig 6A) differs from our experimental data (Fig 2). Therefore, this model still somewhat overestimates the contribution of the adhesion stress to the dynamics of phagocytic spreading. Nevertheless, of the two considered models only the protrusive zipper model succeeds in capturing the essential dynamics of experimentally observed phagocytic spreading, establishing that active cellular protrusion rather than passive cell-substrate adhesion is the dominating driving force of this type of cellular motion.

### Protrusive zipper model with discrete adhesion sites: best match between experiments and theory

The previous sections presented continuum models of cell spreading, resulting in predictions that overestimated the attractive force due to cell-substrate adhesion even in the case where short-range adhesion plays only a secondary role. In reality, however, adhesive ligands encountered by a spreading cell reside at discrete substrate locations rather than presenting a contiguous adhesive film. A pioneering study that considered discrete adhesion sites not only predicted dramatic qualitative differences compared to continuum models of adhesion but even concluded that, depending on the lateral spacing of discrete cell-substrate cross-bridges, “there is little or no tendency for the contact to spread unless the surfaces are forced together” [43]. These considerations prompted us to design and examine a version of the protrusive zipper model in which the model cell can only form new attachments to the substrate at prescribed locations (Fig 7, S5 Movie).

**Fig 7.**
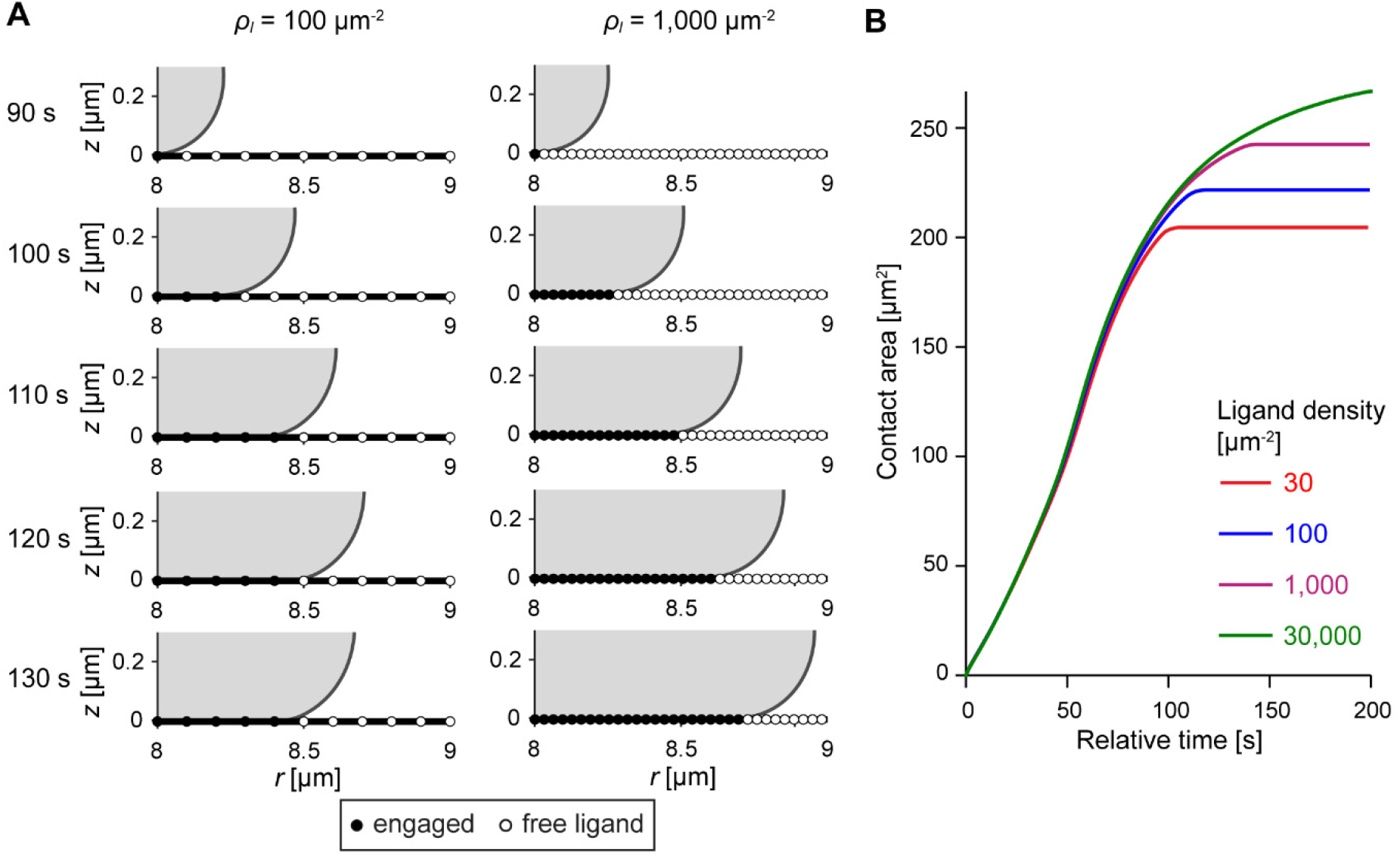
Predictions of the protrusive zipper model with discrete adhesion sites. (A) Two time series (times shown at the left) of simulation snapshots illustrate how the progression of the model cell’s leading edge is affected by the spacing of discrete adhesion sites. Unoccupied binding sites are depicted by empty circles, whereas filled circles indicate that the cell membrane is locally attached. (B) Contact-area-versus-time curves share similar slopes over three orders of magnitude of ligand density, but plateau at different maximum contact areas.

In this model version, the spacing of adhesion sites is determined by the ligand density *ρ_l_* (the number of ligand molecules per μm^2^), with the in-plane spacing given by (*ρ_l_*)^−1/2^. Furthermore, we introduce two additional rules to capture realistic aspects of cell spreading on discrete adhesion sites. First, we replace the continuous adhesion stress (Eq (6)) with the following concept of discrete bond formation. We consider a new bond as “formed” as soon as the distance between the next free ligand site and the closest point of the advancing cell membrane reaches a threshold distance *d_thresh_*. As a consequence of such bonding, we place the cell surface into irreversible contact with this substrate site when carrying out computations at the next time step. The value of the threshold distance is determined by the magnitude of membrane fluctuations. Assuming that the time scale of membrane fluctuations is generally smaller than our time step, we define *d_thresh_* to be the root mean square height *<h^2^>^½^* of membrane fluctuations at equilibrium through [44]:

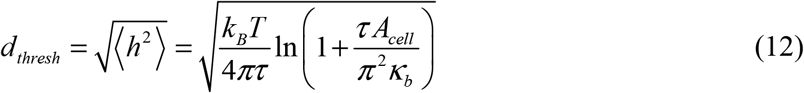

where *k_B_* is Boltzmann’s constant, *T* is temperature, *A_cell_* is the cell surface area, and *κ_b_* is the membrane bending modulus. According to this rule, increased cortical tension *τ* suppresses membrane fluctuations, making it more difficult for the cell to form new adhesive connections at later time points. The parameter values chosen in our simulations (S7 Table) correspond to typical fluctuation amplitudes ranging from 18 nm at resting tension to about 2 nm for very high tension, in general agreement with measurements of membrane fluctuations in cells on surfaces [45, 46].

Second, because it seems unlikely that persistent receptor-ligand contacts induce signaling indefinitely, we also assume that the protrusion stress can diminish over time. This assumption is supported by experimental data showing that actin polymerization does not continue indefinitely during phagocytosis if the cell encounters an unopsonized region of the target particle [23, 47, 48]. We implement this behavior by postulating a transient protrusion stress that initially is triggered by fresh cell-substrate contact (occurring at time *t_bind_*) and then decays exponentially according to the relation:

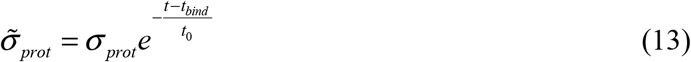

In this version of the model, *σ_prot_* in Eq (8) is replaced by 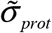 defined in Eq (13). The effects of varying the rate of this decay are explored in S6 Appendices, Appendix F. We use a value of *t_0_* = 66 s in our simulations, which gave the best quantitative agreement with experimental data.

To avoid possible bias due to the exact placement of discrete binding sites, we performed each simulation multiple times, shifting all binding sites laterally by small increments without changing their spacing. We then averaged the results of these individual simulations to predict the cell-spreading behavior as a function of the chosen spacing—or density—of adhesion sites.

Overall, the predictions of this discrete protrusive zipper model (Fig 7) agree well with those of the continuum model version. For instance, maximum cell-substrate contact areas exhibit a similar dependence on the ligand density in both models. However, in this discrete model, the sigmoidal contact-area-versus-time curves obtained at different ligand densities are aligned much more closely during the spreading phase, reflecting spreading speeds that hardly depend on the ligand density anymore (Fig 7B).

Together, the predictions by the model with discrete adhesion sites agree well with the behavior of real human neutrophils during frustrated phagocytosis (Fig. 8). As in experiments, the spreading speed remains remarkably consistent over a range of ligand densities spanning 3 orders of magnitude (Figs 7B and 8A). Moreover, the predicted moderate increase of the maximum contact area at higher ligand densities matches experimental measurements quite well (Fig 8B). On the other hand, whereas the measured dependence of the maximum contact area on the IgG density appears to reach a plateau for IgG densities greater than 1,000 μm^−2^, our model predicts a continuous increase of the maximum contact area over the whole range of ligand densities. A possible explanation for this discrepancy is the model assumption that the ligand density determines the maximum number of discrete adhesion bonds between the cell and the substrate, which implies that the receptors on the cell surface are always in excess. Human neutrophils indeed present a large number of Fcγ receptors on their surface, i.e., several thousand receptors per square micrometer. However, this receptor density is smaller than the highest ligand density of ~25,000 IgG molecules per square micrometer used in our experiments. Therefore, at the highest IgG densities, we expect the numbers of receptors to be the limiting factor determining the maximum number of adhesion sites, which is not accounted for in the above model. To verify this expectation, we conducted additional proof-of-principle simulations in which both the ligands as well as the receptors were treated as discrete, as explained in S6 Appendices, Appendix F. This approach indeed improved the agreement between experiments and theory further, as shown in Fig. 8B.

**Fig 8.**
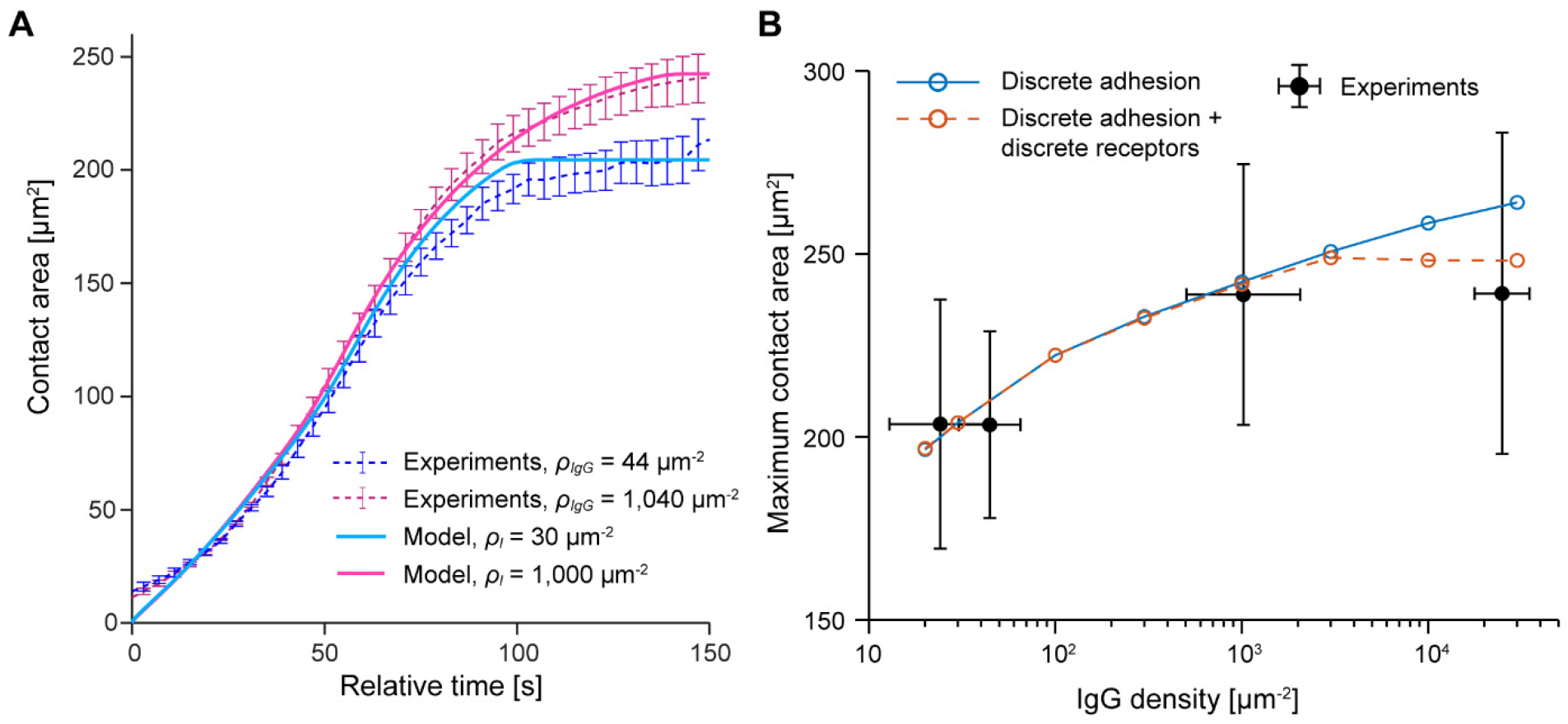
Direct comparison of experimental data with the predictions of the protrusive zipper model with discrete adhesion. (A) Time courses of the cell-substrate contact area predicted by the protrusive zipper model with discrete adhesion are overlaid on experimental results for two different ligand densities. The model curves are not the result of nonlinear fits but simply the predictions based on reasonable choices of parameter values. (B) The predicted ligand-density-dependent increase in maximum contact area agrees well with experimental results obtained in frustrated phagocytosis experiments. Modeling only ligands as discrete entities results in an apparent overestimation of the maximum contact area at the highest IgG density (*solid blue curve*), indicating that the number of Fcγ receptors rather than the number of IgG ligands becomes the limiting factor of the maximum adhesion strength in this regime. A model version that accounts for the discrete nature of both ligands as well as receptors (S6 Appendices, Appendix F) improves the agreement with the data at the highest IgG density (*dashed red curve*). Error bars indicate standard deviation.

Overall, these results reinforce the conclusion that active cytoskeletal protrusion drives phagocytic spreading. They also highlight that it can be important for realistic models of cell spreading on substrates coated with specific adhesion molecules to account for the discrete distribution of these adhesion sites.

## Discussion

Quantitative understanding of the behavior of motile mammalian cells—from immune cells to metastasizing cancer cells—is a prerequisite for transformative biomedical advances in diagnosis, treatment, and therapy of many current and future health threats [49]. Motile cells integrate a complex variety of physical and chemical cues to carry out specific functions. Genuine understanding of the underlying mechano-chemical processes not only requires an interdisciplinary strategy but also integrative experimental/theoretical approaches. Translation of mechanistic notions into mathematical models, and comparison of theoretical predictions with experimental observations, often provides the strongest arguments for or against the validity of biological hypotheses. Confidence in the predictive power of computer simulations requires that the model assumptions be realistic and biologically plausible. Guided by this perspective, we here examined fundamental biophysical mechanisms that potentially can facilitate spreading of biological cells on a substrate.

Unpolarized cells usually begin spreading in a symmetric fashion [8, 50]. This stage is often viewed as a passive process that is entirely driven by adhesion energy due to receptor-ligand binding [8, 51]. An alternative class of models considers such spreading as dependent on cytoskeletal activity [10, 14, 15]. In this study, we have developed a mechanical modeling framework that allows us to establish whether frustrated phagocytosis, a specific form of isotropic cell spreading, is primarily driven by adhesion (Brownian zipper model) or by protrusive forces generated by the cell (protrusive zipper model). Our findings likely carry over to other types of cell spreading inasmuch as key characteristics of adhesion and protrusion are shared [52].

Our results invite a careful review of mathematical models of phagocytosis or other types of cell spreading that consider pre-contact adhesive attraction between cell and substrate as the main driving force [8, 13, 26]. A first important prerequisite for the validity of such models is the use of realistic values of basic mechanical parameters, such as the effective viscosity of the cell interior, the cortical tension, and the adhesion energy density. For example, if the equilibrium cell-substrate contact areas measured in our spreading experiments on low-to-moderate ligand densities were due to adhesion, then the involved adhesion energies would have to be several orders of magnitude higher than those estimated from our measurements of ligand density. Even receptor accumulation in the contact region could not make up for this difference, because the smaller number of available ligands limits the number of possible adhesion bonds. Furthermore, the ability to simulate not only equilibrium situations but also the dynamical cell behavior, and to leverage time-dependent experimental observations against the predictions of such models, considerably raises confidence in a model’s evaluation. As demonstrated in this study, such comparison greatly benefits from the availability of experimental data obtained on substrates presenting a large range of known densities of adhesive ligands.

It appears that adhesion-driven spreading is largely limited to situations where adhesion is unusually strong, and where the cells—like red blood cells—or surrogate objects—like lipid vesicles—are characterized by a low-viscosity interior and very low tension of their cortex or membrane. Indeed, a recent mechanical modeling study found that Brownian membrane fluctuations were only sufficient to drive very small amounts of cell spreading on a surface [53]. If there are cases in which motile mammalian cells spread predominantly due to adhesion, then such spreading will likely occur on much longer timescales than considered here. Finally, it may be possible that adhesion plays a more relevant role in the engulfment of very small particles, or very early during phagocytic spreading, where cell shape changes do not require a significant expansion of the cell surface area [54]. Irrespective of the relevance of Brownian-zipper-type models in a particular practical situation, we note that the excellent agreement of our theoretical treatment of purely adhesion-driven spreading with other studies focusing on this form of motion [8, 13] strengthens confidence in our modeling approach and its numerical implementation.

The only alternative mechanism by which cells can expand their region of contact with a suitable substrate appears to be active, protrusion-dominated spreading. Many imaging studies indeed have documented an increased density of F-actin at the protruding front of motile immune cells [16, 55, 56], supporting the biological plausibility of the protrusive zipper concept. Moreover, inhibition of actin polymerization with cytochalasin B or latrunculin A eliminates the rapid phase of fibroblast spreading [10, 14, 57]. Similarly, chemical disruption of the actin cytoskeleton generally inhibits phagocytosis by immune cells [36, 47]. We have previously shown that extremely small concentrations of actin inhibitors actually can increase the size of transient, chemotactic-like protrusions [36, 58]. However, the cortical tension was reduced in such cases, and phagocytic spreading was not enhanced, highlighting the multiplicity of—likely dichotomous—mechanical roles of F-actin during cell deformations.

Our simulations based on the protrusive zipper model not only reproduce the main features of neutrophil spreading on IgG-coated substrates but, in the case of discrete adhesion sites, they also agree very well with experimental data. We note that this agreement neither is surprising, nor does it constitute solid proof of the validity of this model. The scarcity of quantitative insights into the underlying mechanisms simply does not place sufficient constraints on the design of computational models. Instead, our goal here was to explore what set of minimalistic rules would need to be postulated to achieve a good match between theory and experiment. The successful implementation of this approach allows us to propose likely mechanistic details that can then provide guidance for future studies. Still, we are confident that the protrusive zipper model paints a reasonably accurate picture of phagocytic spreading, not only because of the elimination of the only alternative hypothesis, but also because our semiempirical model is plausible, i.e., it does not contradict pertinent biophysical and biological insights. For example, our implementation of the relationship between receptor-induced signaling and the generation of protrusive force is consistent with a successful previous model of phagocytosis [22], as well as with previous measurements of the biphasic signaling response from a macrophage-like cell line during Fcγ receptor-mediated phagocytosis [42]. Models based on the assumption that actin polymerization preferentially occurs in regions of higher membrane curvature [59] can result in a localization of protrusion stress that is similar to the form it takes in our model (Fig 1D), although the idea of curvature-induced actin polymerization seems problematic in view of the much higher membrane curvatures in ever-present surface wrinkles and microvilli of immune cells. Importantly, our implementation of the relationship between receptor-induced signaling and force generation is independent of the exact mechanism by which intracellular processes are translated into protrusive force; instead, it encompasses different mechanistic notions such as the Brownian ratchet [60] and end-tracking motors [61].

An important outcome of our simulations of the protrusive zipper model is the difference in predictions depending on whether we considered continuous or discrete cell-substrate adhesion, in agreement with a previous theoretical study [43]. We are unaware of physiological situations in which cell adhesion is not predominantly supported by specific receptor-ligand bonds. Thus, the model version that is based on discrete binding sites is the more realistic version of the protrusive zipper. Remarkably, it is also the version that yielded the best match between theory and experimental observations. To achieve this agreement, we postulated that the formation of new receptor-ligand bonds is coupled to the protrusion stress in a time-dependent manner (Eq (13)), based on the assumption that receptor-induced signaling and subsequential actin polymerization are not sustained indefinitely after a ligand activates the receptor. There are multiple factors that could lead to this effective decay, such as the depletion of precursor molecules needed for the production of signaling messengers (e.g. release of IP3 will cease once PIP2 has been locally depleted [62]).

Perhaps the most important insight from the comparison of model versions based on continuous versus discrete adhesion is that models based on continuous adhesion are likely to misrepresent key aspects of the cell behavior, especially on lower ligand densities. For example, such models cannot capture the possibility that too large ligand spacing may prevent the next unbound receptor-ligand pair from approaching one another to within the distance at which they are likely to form a bond. In our protrusive zipper model with discrete adhesion, variation of substrate adhesivity by changing the density of adhesion sites primarily affects the maximum cell-substrate contact area. This prediction agrees well with our experiments, unlike the predictions by model versions that treat the ligand layer as a continuum.

The mechanisms governing isotropic cell spreading studied here are likely to also play a crucial role in the directional motion of migrating cells. Many immune cells, for example, are capable of executing both types of motion, as required during phagocytic spreading versus chemotactic and other forms of migration. As revealed by our experiments and reproduced by our discrete adhesion model, the rate of isotropic cell spreading is largely conserved on substrates presenting a broad range of densities of adhesion sites. This finding exposes an interesting difference to the prediction by models of migrating cells, i.e., that the crawling speed of such cells has a maximum on substrates with intermediate adhesion strengths [63–65]. This difference is a reminder that directional migration generally is a more complex type of motion that involves, in addition to protrusion and adhesion, important mechanisms regulating contraction and detachment of the rear of the migrating cell from the substrate. The positive correlation between low-to-moderate ligand densities and the speed of motion predicted by models of migrating cells can be attributed to either a direct scaling of protrusion with the amount of adhesion [64], or a postulated increased actin retrograde flow at lower adhesion strengths [63]. The predicted slowing of migrating cells at higher ligand densities is caused by an effective frictional force due to the need to break stronger adhesive attachments at the cell’s rear [66, 67]. A noteworthy recent model of cell migration that decoupled friction from adhesive attraction actually predicted faster movement for higher adhesion energies [51]. The excellent agreement between our experiments and the protrusive zipper model suggests that the above additional mechanisms included in models of migrating cells play at most a minor role in isotropic phagocytic spreading. On the other hand, we believe that the main insights exposed by our study provide important suggestions on how to treat the interplay between protrusion and adhesion in models of migrating cells.

## Methods

### The perturbed Stokes equations

Assuming the cell interior can be approximated by a single, highly viscous, slow-moving Newtonian fluid, its motion obeys the Stokes equations for creeping flow with no body force, as given by Eq (2). Furthermore, for an incompressible fluid, the equation of continuity simplifies to the incompressibility condition (Eq (3)). Due to axial symmetry, there are three unknowns: *v_r_, v_z_*, and *p*, where *r* and *z* are cylindrical polar coordinates.

To solve this system of partial differential equations, we use the finite element method (FEM). Because directly solving the Stokes equations using FEM leads to numerical instabilities [68], we follow the procedure outlined by Drury and Dembo [37] to solve a perturbed form of the Stokes equations, in which the incompressibility condition (Eq (3)) is approximated as

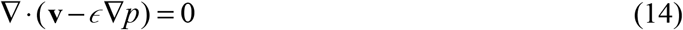

In this form, the fluid is nearly incompressible, which allows us to avoid the well-known problems with using FEM for incompressible problems [68]. To accurately solve the Stokes equations, the value for *ϵ* must be sufficiently small. Specifically, we use the condition given in [37]:

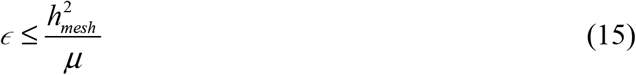

where *h_mesh_* is the characteristic radius of a single quadrilateral element. We tested different values of *ϵ* in our models and found this condition to provide sufficient accuracy and convergence (S6 Appendices, Appendix E).

Over the unbound portion of the cell, we specify Neumann boundary conditions, corresponding to a stress balance (Eq (4)). For the adherent cell surface, we specify Dirichlet boundary conditions, simply a no-slip condition in this case (Eq (9)). The pressure boundary condition is given by assuming the normal component of the pressure gradient goes to zero at the cell surface:

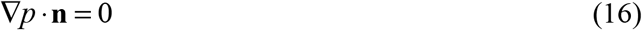

### Computational details

At each time step, we first compute values for the surface curvature of the cell. Local values of the angle of the normal *ϕ* are evaluated using cubic polynomial fits. We then calculate the boundary stresses using the relationships outlined in the main text. The integral for the effective adhesion stress (Eq (6)) is computed numerically as described in S6 Appendices. The values for tension and protrusion stress at each time step are evaluated using smoothed versions of the relationships given in Eq (1) and Eq (11) (see S6 Appendices, Appendix C for full forms).

To find the fluid velocity and pressure fields, we must simultaneously solve Eq (2) and Eq (14) for *v_r_, v_z_*, and *p*. We employ the strategy outlined in [37], which is an adaptation of the Uzawa method. Starting with an estimate for pressure *p_est_* given either by the previous time step or as a uniform distribution prior to spreading, we solve for fluid velocities by treating the pressure as known in Eq (2):

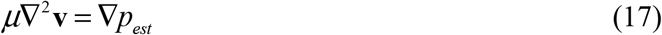

We then treat the fluid velocities as known (**v**_*est*_) and solve an altered version of equation (14):

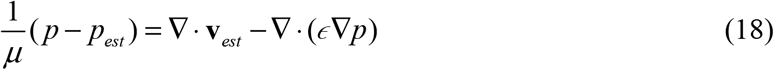

The updated pressure is then inserted back into Eq (17) and the procedure is repeated until the following condition is met:

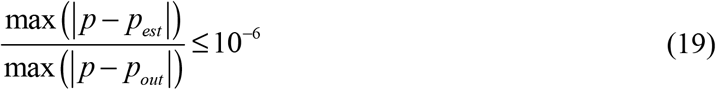

This generally occurs in less than 50 iterations.

The new cell boundary is then given by

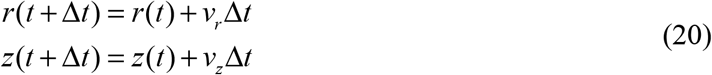

The value for Δ*t* was set relative to the Courant number for each iteration, such that fluid flow occurs over at most 10% of any individual element (S6 Appendices). In the continuum adhesion case (no discrete adhesion sites), the contact line is updated at each time step by computing the intersection between the cell boundary and the surface, then adding all points advected to *z* ≤0 to the Dirichlet boundary condition (contact, no-slip) for the next time step. In the discrete adhesion model, contact is determined by the distance threshold computed from Eq (12). The contour is smoothed after each advection, and then small corrections are made to the cell contour to compensate for any deviations from constant volume due to smoothing and numerical error.

This model was implemented in MATLAB R2020b (MathWorks) and the full code is available at https://github.com/emmetfrancis/phagocyticSpreading. Individual simulations generally run within 1-3 hours on a personal computer.

## Supporting information

S7 Table (parameter values)

## Acknowledgements

We thank Dr. Robert Guy from UC Davis Mathematics for many helpful conversations on various iterations of our model and for assistance with the derivation found in S6 Appendices, Appendix B. We also thank Dr. Marc Herant for fruitful discussions on computational modeling of motile cells and Dr. Vasilios Morikis for his feedback on early versions of this manuscript.

## Supporting Information

**S1 Movie. Simulations of purely adhesive cell spreading (“Brownian zipper”) on substrates presenting different densities of adhesion sites.** This sequence of animations shows side views of model cells spreading on adhesive substrates coated with relative ligand densities (*ρ_l_*) of 1%, 3%, 10%, 30%, and 100% for 400 s in the absence of protrusion stress.

**S2 Movie. Simulation of purely protrusion-driven cell spreading.** This animation shows a side view of a model cell spreading on a flat substrate for 200 s. The cell motion is assumed to be driven exclusively by active cytoskeletal protrusion at a maximum protrusion stress of *σ_prot,max_* = 3.5 kPa. Cell-substrate contact, once established, is assumed to be permanent. All other contributions of adhesive cell-substrate interactions to spreading are neglected.

**S3 Movie. Effect of the range of action of the protrusion stress on purely protrusion-driven cell spreading.** This sequence of simulations shows side views of model cells spreading in the absence of adhesion stress for 200 s. Increasing the range *s*_0_ over which cytoskeletal protrusion actively pushes the cellular pseudopod outwards (given in terms of the arc length of the free cell contour, measured from the point of substrate contact) results in thicker pseudopods. In this sequence, *s_0_* = 0.4 μm, 1.2 μm, and 2.0 μm, respectively. In all examples, *σ_prot,max_* × *s_0_* = 2.8 mN/m.

**S4 Movie. Minor role of cell-substrate adhesion stress in protrusion-dominated cell spreading (“protrusive zipper”).** This sequence of simulations compares magnified views of purely protrusion-driven spreading (adhesion stress *ρ_l_* = 0%) with protrusion-driven spreading in the presence of moderate and strong adhesion stress (*ρ_l_* = 10% and 100%, respectively). In all examples, *σ_prot,max_* = 3.5 kPa.

**S5 Movie. Protrusive zipper simulations with discrete adhesion sites.** This sequence of simulations shows a magnified view of protrusion-driven spreading (*σ_prot, max_* = 3.5 kPa) on substrates presenting different densities of discrete adhesion sites (*ρ_l_* = 30 μm^−2^, 100 μm^−2^, 1,000 μm^−2^, and 10,000 μm^−2^). Open circles denote free adhesive binding sites, and filled circles depict sites with engaged ligand.

**S6 Appendices. Derivation of important relationships and summary of numerical testing.** (A) Adhesion stress calculation. (B) Derivation of a power law for contact area growth in passive spreading. (C) Development and testing of constitutive relations for cortical tension and protrusion stress. (D) Details of finite element implementation. (E) Summary of numerical testing. (F) Additional testing of the discrete adhesion model.

**S7 Table. Summary of modeling parameters, with supporting references.**

