## Supplementary material for "Integrative experimental/computational approach establishes active cellular protrusion as the primary driving force of phagocytic spreading by immune cells": S6 Appendices

### Appendix A. Adhesion stress calculation

To compute the adhesion stress, we consider a potential acting between an adhesion molecule and a receptor in the cell membrane. Here, we use a 6-3 potential:

$$u_{adh}(D) = u_0\left[\left(\frac{D_0}{D}\right)^6 - 2\left(\frac{D_0}{D}\right)^3\right] \tag{S1}$$

where $D_0$ is the distance associated with the energy minimum and $u_0$ sets the magnitude of the binding energy. The force due to this single interaction is

$$f_{adh}(D) = -\frac{du_{adh}(D)}{dD} = f_0\left[\left(\frac{D_0}{D}\right)^7 - \left(\frac{D_0}{D}\right)^4\right] \tag{S2}$$

where $f_0 = 6u_0/D_0$. As $D_0$ is the location of the energy minimum, we also see that $f_{adh}(D_0) = 0$. We use $D_0 = 50$ nm in our simulations. Realistically, one might expect this distance to be much smaller (molecular scale), but the longer range here effectively includes membrane fluctuations and filopodia and serves as an upper limit. Adhesion potentials with a range of tens to hundreds of nanometers are common when modeling cells and vesicles at the mesoscopic scale [1, 2].

To solve for the overall adhesion stress, we integrate over the ligand-coated surface (surface coordinates $[r, \theta]$ ) and the region of membrane being considered (surface coordinates $[s, \psi]$ ) (Fig S1A).

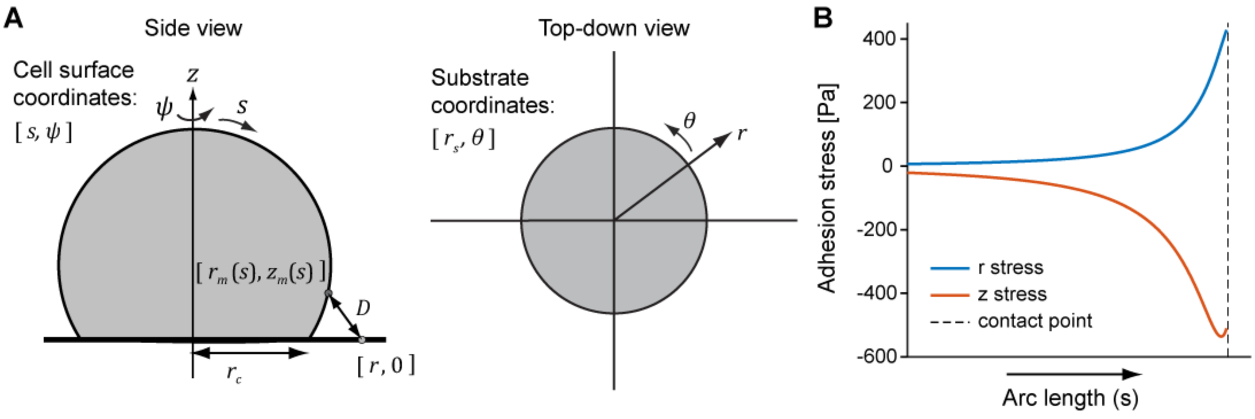


**Fig S1. Adhesion force calculation.**
(A) The cell geometry is depicted from the side, as well as from the top, including definitions of coordinates used in the text. (B) Adhesion stress components are plotted as a function of arc length $s$ for the cell shape shown in part A at a relative ligand density of 10%.

Individual interaction forces are specified as vectors, that is

$$\mathbf{f}_{single} = f_0 \left[ \left( \frac{D_0}{D} \right)^7 - \left( \frac{D_0}{D} \right)^4 \right] \mathbf{f}_{unit} \tag{S3}$$

The distance between a point on the membrane and a point on the surface is given by

$$D(r,\theta,s,\psi) = \sqrt{(r_m \cos\psi - r\cos\theta)^2 + (r_m \sin\psi - r\sin\theta)^2 + z_m{}^2} \tag{S4}$$

The distance $z_m$ is adjusted from the $z$ coordinate used elsewhere so that the surface $z = 0$ corresponds to the zero-force distance $D_0$, i.e. $z_m = z + D_0$.

The direction of the force is expressed by a unit vector (we only calculate the $r$ and $z$ components, as the angular component cancels out due to symmetry)

$$\mathbf{f}_{unit} = \frac{1}{D} \begin{bmatrix} r_m \cos\psi - r\cos\theta \\ z_m \end{bmatrix} \tag{S5}$$

Considering a region of the cell membrane extending from $s = s_{cur} - l/2$ to $s = s_{cur} + l/2$ along the arc length and from $\psi = -\Delta\psi/2$ to $\psi = \Delta\psi/2$ in the angular direction, we compute the pairwise force by the total integral

$$\mathbf{f}_{adh} = \int_{-\frac{\Delta\psi}{2}}^{\frac{\Delta\psi}{2}} \int_{s_{cur}-\frac{l}{2}}^{s_{cur}+\frac{l}{2}} \left( \int_0^{2\pi} \int_{r_c}^{\infty} \mathbf{f}_{single} \rho_l r dr d\theta \right) \rho_R r_m ds d\psi \tag{S6}$$

where $\rho_l$ is the ligand density on the substrate and $\rho_R$ is the receptor density on the cell membrane. In practice, we lump $f_0$ and $\rho_R$ into a single parameter $\sigma_0 = f_0\rho_R$ , which effectively serves as a force per unit area of cell membrane (see S7 Table).

The stress per unit area of membrane is given by

$$\boldsymbol{\sigma}_{adh} = \frac{\mathbf{f}_{adh}}{\int_{-\frac{\Delta\psi}{2}}^{\frac{\Delta\psi}{2}} \int_{s_{cur}-\frac{l}{2}}^{s_{cur}+\frac{l}{2}} r_m ds d\psi} \tag{S7}$$

In the limit $l\Delta\psi \rightarrow 0$ , we obtain the final expression, as given in the main text

$$\boldsymbol{\sigma}_{adh} = \sigma_0 \rho_l \int_0^{2\pi} \int_{r_c}^{\infty} \mathbf{f}_{unit} \left[ \left( \frac{D_0}{D} \right)^7 - \left( \frac{D_0}{D} \right)^4 \right] r dr d\theta \tag{S8}$$

This integral is not readily carried out analytically. Therefore, we evaluate it numerically in MATLAB by performing trapezoidal integration in both dimensions, using log-spaced meshes for both $r$ and $\theta$. Results of this integration are essentially identical to those of integration with MATLAB built-in functions, but computation time is reduced well over 10-fold. An example of the adhesion stress components computed as a function of arc length $s$ along the cell contour defined in Fig S1A is shown in Fig S1B for $\rho_l = 10\%$.

In practice, we modify the adhesion strength in simulations by varying the scaling factor $\sigma_0\rho_l$ , but we cannot directly extract the ligand density $\rho_l$. Instead, we infer the adhesion energy per unit area $\gamma$ from the equilibrium contact area $A_c$ using simple geometry, conservation of volume, and the Young-Dupre equation, as explained in the main text. Using these estimated values of adhesion energy $\gamma(A_c)$ and the typical binding energy of a low affinity Fcγ receptor to IgG ($E_{bind} \approx 6 \times 10^{-20}$ J [3]), we estimate the IgG density $\rho_{IgG,est}$ as

$$\rho_{IgG,est} = \frac{\gamma(A_c)}{E_{bind}} \quad \text{(S9)}$$

We perform a linear fit between $\rho_{IgG,est}$ and the relative adhesion strength $\rho_l / \rho_{l,max}$ to compute the effective IgG densities tested in our Brownian Zipper model (Fig S2B). From this fit, we find that $\rho_l = 10\%$ roughly corresponds to $\rho_{IgG,est} = 1{,}000\ \mu m^{-2}$.

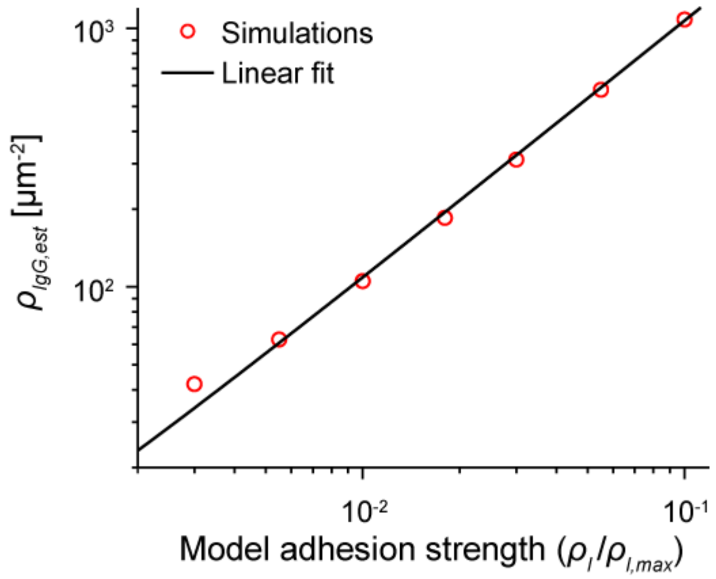


**Fig S2. Relationship between estimated IgG density and model adhesion strength**.
A linear fit relates model adhesion strength to IgG density based on the equilibrium contact areas obtained from simulations.

### Appendix B. Derivation of a power law for contact area growth in passive spreading

To develop an approximate analytical prediction of the functional dependence of the contact area on time, we start from the overall energy balance for our passive model cell spreading on a flat surface with adhesion energy density $\gamma$, analogous to the energy balance in the spreading of a viscous droplet [4]. After canceling a factor of 2π, this yields

$$\frac{\gamma}{2}\frac{\partial}{\partial t}(r_c^{\,2}) = 2\mu\iint e_{ij}e_{ij}rdrdz + \frac{\partial}{\partial t}\left(\tau\int rds\right) + \frac{\partial}{\partial t}\left(\iint prdrdz\right) \tag{S10}$$

where $r_c$ is the cell contact radius, $\mu$ is the effective cell viscosity, $e_{ij}$ denotes the $i,j$ component of the rate-of-strain tensor (repeated indices are summed over), $\tau$ is the cortical tension, and $p$ is the pressure inside the cell. The left-hand side corresponds to energy gain due to cell-surface adhesion. The first term on the right-hand side is the viscous dissipation rate [5], the second term is the work required to deform the cortex, and the third is the change in energy due to changes in pressure and/or volume. In our simulations volume is kept constant, and any energy required for pressure changes is much lower than for changes of the surface area or tension; therefore, we neglect the last term of Eq (S10) in the following analysis.

To estimate the remaining terms in the energy balance, we assume that the passively spreading cell adopts the shape of a spherical cap with contact radius $r_c$ and radius of curvature $R_{cell}$ (Fig S3A). This geometry defines the contact angle as:

$$\sin\theta_c = \frac{r_c}{R_{cell}} \tag{S11}$$

We focus this analysis on initial contact area growth, during which the contact angle is small and we can treat $R_{cell}$ as a constant. Based on this geometry and features of the calculated flow profile (Fig S3A inset), we estimate the viscous dissipation rate and work required to deform the cortex as follows.

We first expand the dissipation term to include all non-zero components of the rate-of-strain tensor (all derivatives with respect to the polar angle are zero given axial symmetry):

$$\Pi = 2\mu\iint e_{ij}e_{ij}rdrdz = \mu\iint\left[2\left(\frac{\partial v_r}{\partial r}\right)^2 + 2\left(\frac{v_r}{r}\right)^2 + 2\left(\frac{\partial v_z}{\partial z}\right)^2 + \left(\frac{\partial v_r}{\partial z} + \frac{\partial v_z}{\partial r}\right)^2\right]rdrdz \tag{S12}$$

The main effect of the adhesion force is to "pull down" the membrane onto the flat surface; that is, the $z$-component of the adhesion stress generally dominates (Fig 1B). Accordingly, we expect $v_z$ to dominate over $v_r$ in the portion of the cell closest to the substrate, which is confirmed by our calculations (Fig S3A inset). Furthermore, due to incompressibility, we know

$$\frac{v_r}{r} + \frac{\partial v_r}{\partial r} + \frac{\partial v_z}{\partial z} = 0 \tag{S13}$$

Therefore, if the contributions of $v_r$ are small, $\partial v_z/\partial z$ must also be small, and we are left with:

$$\Pi \approx \mu \iint \left( \frac{\partial v_z}{\partial r} \right)^2 r dr dz \tag{S14}$$

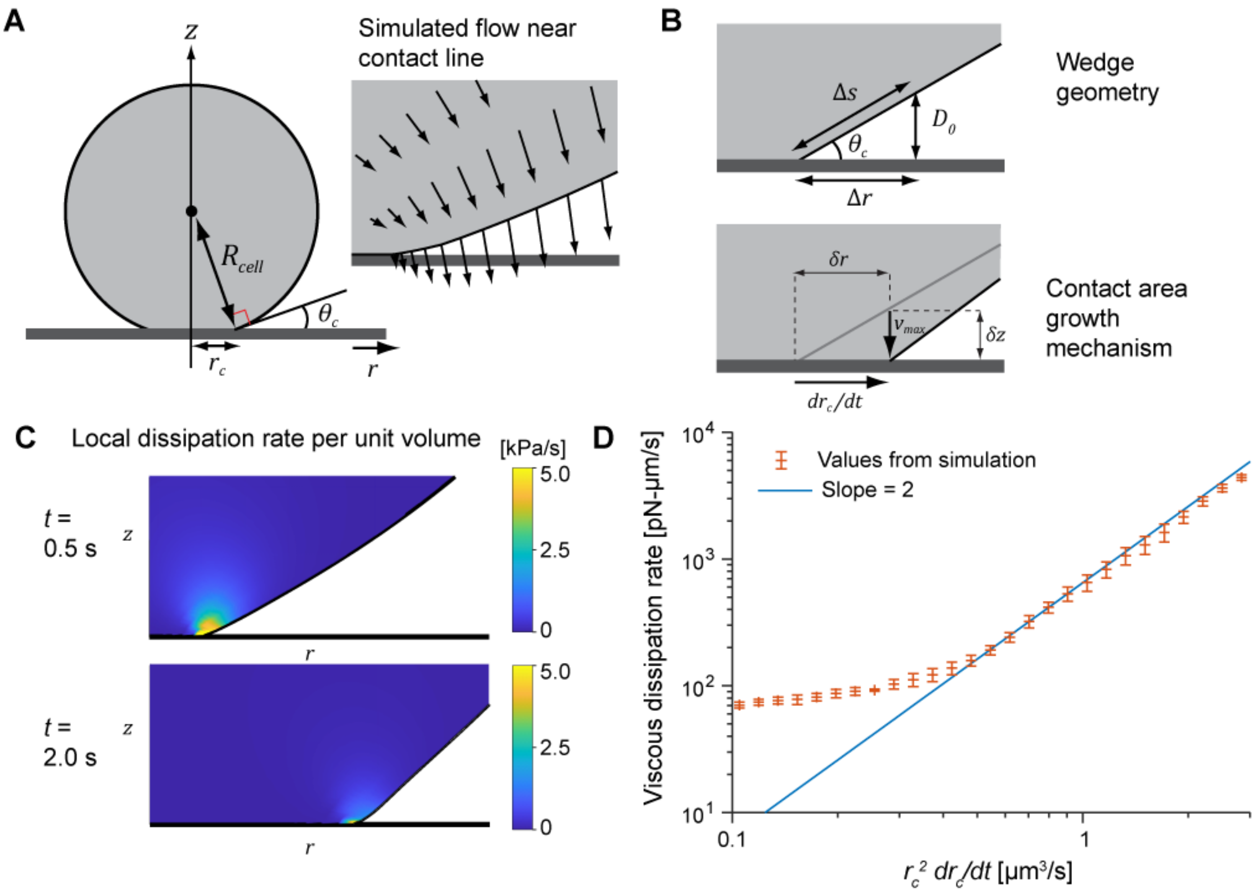


**Fig S3.** Energy balance calculations.
(A) Geometry of a spreading droplet during early spreading. The inset illustrates fluid velocities (arrows) at the boundary near the contact line as predicted by a typical Brownian zipper simulation. (B) Simplified geometry of a wedge used to represent a spreading droplet. The top panel defines variable used in the text. The bottom panel illustrates the manner of contact area growth implemented in our model. (C) Local dissipation per unit volume at the contact line for two early time points as computed by a Brownian zipper simulation at a ligand density of 10%. The dissipation is tightly confined to the point of contact. (D) Total viscous dissipation rate was directly computed using linear shape functions and plotted against $r_c^2\ dr_c/dt$ on a log-log axis. Simulated values were binned and averaged for each interval on the x-axis. Error bars indicate standard deviation of binned data.

As a test geometry for points near the surface, we consider a wedge-like shape, as shown in Fig S3B. The exact functional form of $v_z$ within this wedge cannot be readily inferred *a priori*. Instead, we identify characteristic length scales to estimate the velocity gradient. We expect the *z*-velocity to be highest close to the surface, that is, within the length scale set by the adhesion potential ($D_0$). The associated distance in the *r*-direction $\Delta r$ (Fig S3B) is given by:

$$\Delta r = \frac{D_0}{\tan\theta_c} \approx \frac{D_0 R_{cell}}{r_c} \tag{S15}$$

Before estimating the velocity gradient, we relate $v_z$ to the rate of contact radius growth, $dr_c/dt$. Because there is no slip at the contact line itself, the only way for the contact area to increase is for nonadherent membrane to approach and touch the surface past the contact line. We consider a sample point close to the surface at which the membrane moves at maximum velocity $v_{max}$ in the negative $z$ direction (Fig S3B). If this point moves down from a distance $\delta z$ from the surface, then the contact line moves outward by a distance of $\delta r$ in the $r$-direction (Fig S3B), given by

$$\delta r = \frac{\delta z}{\tan\theta_c} \tag{S16}$$

In terms of velocities, this yields

$$\frac{dr_c}{dt} = \frac{v_{max}}{\tan\theta_c} \approx \frac{v_{max} R_{cell}}{r_c} \tag{S17}$$

Combining Eqs (S15) and (S17) and assuming the velocity gradient is highest in the direction tangent to the membrane, we approximate the sought velocity gradient:

$$\frac{\partial v_z}{\partial r} \approx \frac{v_{max}}{\Delta r} = \frac{r_c^2}{R_{cell}^2 D_0} \frac{dr_c}{dt} \tag{S18}$$

We insert these estimates into the dissipation term (Eq (S14)) and integrate over the wedge:

$$\begin{aligned} \Pi &\approx \mu \left( \frac{r_c^2}{R_{cell}^2 D_0} \frac{dr_c}{dt} \right)^2 \int_{D_0 \frac{r-r_c}{\Delta r}}^{D_0} \left( \int_{r_c}^{r_c+\Delta r} r dr \right) dz \\ &= \mu \left( \frac{r_c^2}{R_{cell}^2 D_0} \frac{dr_c}{dt} \right)^2 \frac{D_0 \Delta r}{6} \left( 3r_c + \Delta r \right) \end{aligned} \tag{S19}$$

Inserting the definition of $\Delta r$ from Eq (S15) and assuming $\Delta r << 3r_c$, we find

$$\Pi \approx \Pi_0 r_c^4 \left( \frac{dr_c}{dt} \right)^2 \tag{S20}$$

where $\Pi_0 = \mu/(2R_{cell}^3)$. We verify this relationship by comparing it to numerically computed dissipation rates obtained in Brownian zipper simulations (Fig S3C-D). The dissipation rate per unit volume is highest in a small region near the contact line (Fig S3C). We integrate to compute the total dissipation rate at each time step, and then plot this as a function of $r_c^2(dr_c/dt)$ to compare it to our approximate analytical prediction from Eq (S20) (Fig S3D). When plotted on a log-log axis, this curve shows a linear region with a slope of 2, in agreement with the scaling in Eq (S20).

We next estimate the work required to deform the cortex, as given by the second term on the right-hand side of Eq (S10). For a spherical cap of contact radius $r_c$ and height $h$:

$$\frac{\partial}{\partial t}\left(\tau\int rds\right)=\frac{\partial}{\partial t}\left[\tau\left(r_c^{\ 2}+\frac{h^2}{2}\right)\right] \tag{S21}$$

Early in spreading, $h$ is approximately constant and tension changes slowly, therefore

$$\frac{\partial}{\partial t}\left[\tau\left(r_c^{\ 2}+\frac{h^2}{2}\right)\right]\approx 2\tau r_c\frac{dr_c}{dt} \tag{S22}$$

Inserting the approximations from Eq (S20) and Eq (S22) into the energy balance (Eq (S10)) and simplifying, we arrive at the following differential equation:

$$r_c^{\ 3}\left(\frac{dr_c}{dt}\right)=\frac{2(\gamma-\tau)}{\Pi_0} \tag{S23}$$

In early spreading, tension changes more slowly than $r_c$; therefore, it can be treated as a constant. This leads to the spreading law observed in our model as well as in a previous computational model of passive cell spreading [2]

$$r_c\propto t^{1/4}\rightarrow A_c\propto t^{1/2} \tag{S24}$$

### Appendix C. Development and testing of constitutive relations for cortical tension and protrusion stress

To best match the real-life behavior of cells, we developed versions of the piecewise relationships for tension (Eq 1) and protrusion stress (Eq 11) that have continuous first derivatives ($C^1$). We accomplished this by using polynomials to transition between each piecewise domain from the original $C^0$ relations. For tension, we use

$$\tau(A_{cell})=\begin{cases} 0.01+0.16\left(\dfrac{A_{cell}}{A_{cell,0}}-1\right)\ \dfrac{\text{mN}}{\text{m}} & \text{if}\quad \dfrac{A_{cell}}{A_{cell,0}}\le 1.13\\ 0.7405\left(\dfrac{A_{cell}}{A_{cell,0}}\right)^2-1.5135\left(\dfrac{A_{cell}}{A_{cell,0}}\right)+0.7955\ \dfrac{\text{mN}}{\text{m}} & \text{if}\quad 1.13<\dfrac{A_{cell}}{A_{cell,0}}\le 1.39\\ 0.0516+0.545\left(\dfrac{A_{cell}}{A_{cell,0}}-1.26\right)\ \dfrac{\text{mN}}{\text{m}} & \text{if}\quad \dfrac{A_{cell}}{A_{cell,0}}>1.39\end{cases} \tag{S25}$$

The $C^1$ version of protrusion stress requires two separate transition regions, which we achieve using cubic polynomials:

$$\sigma_{prot} = \sigma_{prot,max} \begin{cases} 0.05 + 0.3\left(\dfrac{A_c}{A_{c,trans}}\right) & \text{if} \quad \dfrac{A_{cell}}{A_{cell,0}} \leq 1.13 \\ c_3 A_c^{\,3} + c_2 A_c^{\,2} + c_1 A_c + c_0 & \text{if} \quad 1.13 < \dfrac{A_{cell}}{A_{cell,0}} \leq 1.39 \\ 0.35 + 0.65\left(\dfrac{A_c - A_{c,trans}}{120\ \mu\text{m}^2}\right) & \text{if} \quad \dfrac{A_{cell}}{A_{cell,0}} > 1.39\ \&\ A_c \leq A_{c,trans} + 100\ \mu\text{m}^2 \\ d_3 A_c^{\,3} + d_2 A_c^{\,2} + d_1 A_c + d_0 & \text{if} \quad A_{c,trans} + 100\ \mu\text{m}^2 < A_c \leq A_{c,trans} + 140\ \mu\text{m}^2 \\ 1 & \text{if} \quad A_c > A_{c,trans} + 140\ \mu\text{m}^2 \end{cases} \tag{S26}$$

The $c$ and $d$ coefficients are found by solving the appropriate linear systems. Their values generally depend on the contact area $A_c$ at different values of surface area deformation; namely, at $A_{cell} = 1.13\ A_{cell,0}$ , $A_{cell} = 1.26\ A_{cell,0}$ , and $A_{cell} = 1.39\ A_{cell,0}$. These are derived from sample runs of the simulation.

A comparison of the non-smooth ($C^0$) relationships with their smooth ($C^1$) counterparts reveals only minor differences in the respective graphs (Figs S4A and S5A). Furthermore, simulations performed with $C^1$ vs. $C^0$ relationships result only in slight changes in spreading dynamics that do not affect the main takeaways of this study (Figs S4B-C and S5B).

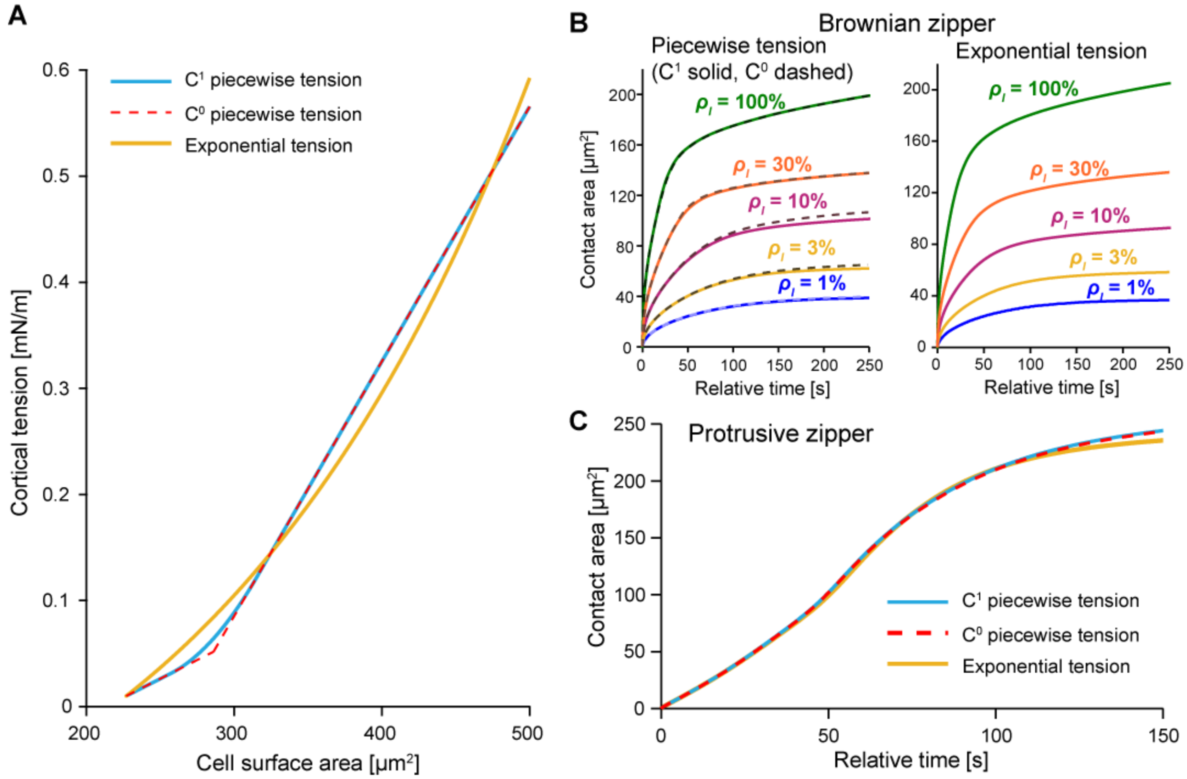


**Fig S4. Results of simulations using alternative cortical tension constitutive relations.**
(A) Alternative relationships between cortical tension and cell surface area. (B) Brownian zipper results do not change qualitatively for different choices of cortical tension constitutive relations. (C) Contact area over time for the pure protrusive zipper varies only slightly between simulations using different relationships for cortical tension.

To confirm that our findings were not dependent on the relationships chosen for tension and protrusion stress, we also tested alternative expressions. For tension, a simpler relationship that has been used in previous numerical studies of cell deformations [6] is exponential:

$$\tau = \tau_0 + 0.15\left( e^{\frac{A_{cell} - A_{cell,0}}{A_{cell,0}}} - 1 \right) \tag{S27}$$

As shown in Fig S4B-C, this relationship does not qualitatively alter the Brownian Zipper or protrusive zipper contact area curves.

A simpler assumption for growth of the protrusion stress is that the stress grows linearly as a function of contact area:

$$\sigma_{prot} = \sigma_{prot,max} \begin{cases} 0.02 + 0.98\left( \dfrac{A_c}{250\ \mu\text{m}^2} \right) & \text{if} \quad A_c \le 250\ \mu\text{m}^2 \\ 1 & \text{if} \quad A_c > 250\ \mu\text{m}^2 \end{cases} \tag{S28}$$

Remarkably, we find that even this simple relationship gives rise to the sigmoidal growth of contact area over time observed experimentally (Fig S5C), indicating this is a general feature of our model that does not depend sensitively on the exact relationship chosen for protrusion stress.

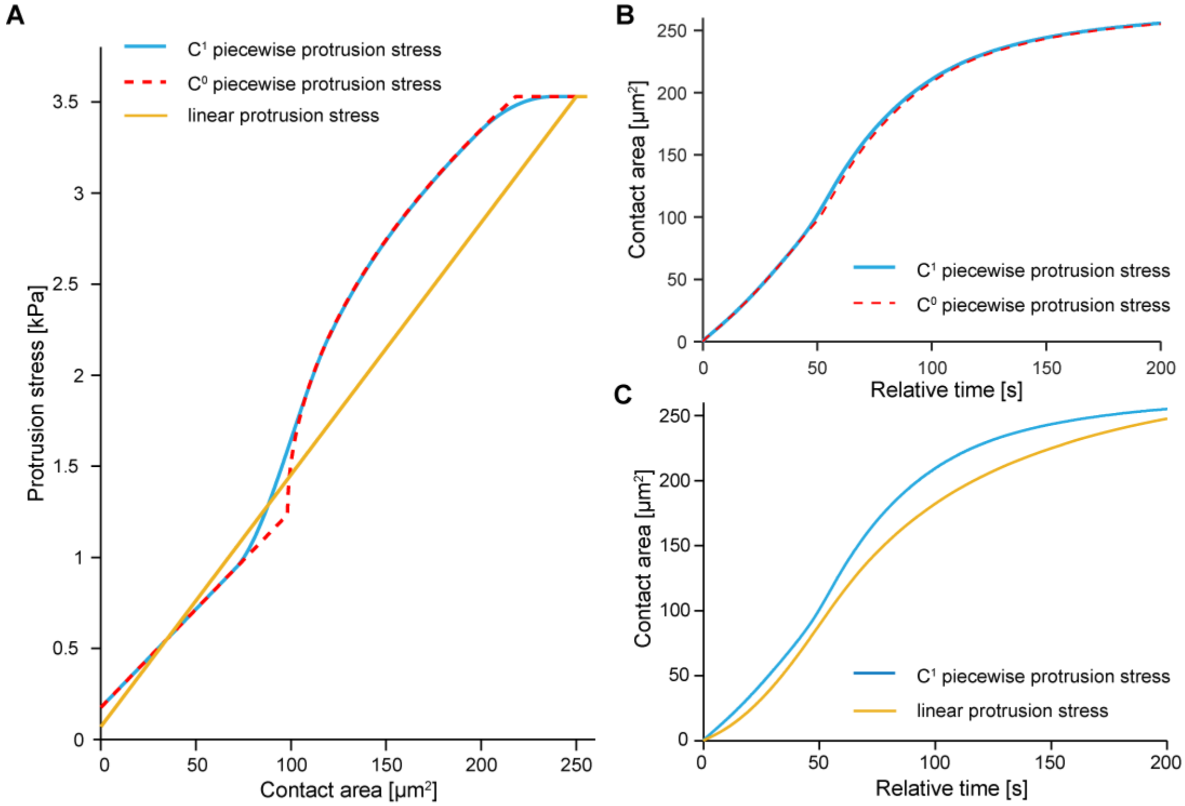


**Fig S5. Results of simulations using alternative protrusion stress constitutive relations.**
(A) Alternative relationships between protrusion stress and cell-substrate contact area. (B) Results of the pure protrusive zipper simulation using the $C^1$ relationship only differ slightly from the $C^0$ version. (C) A linear relationship between protrusion stress and contact area gives rise to a sigmoidal contact area vs. time curve.

### Appendix D. Details of finite element implementation

The governing equations are given in Methods, referred to as the "velocity equation" (Eq 17) and the "pressure equation" (Eq 18) below. Both equations are readily cast into a Galerkin form, which then is rewritten in matrix form as is conventional for finite element methods.

For reference, we give the elemental contributions to the stiffness matrices and load vectors for each finite element system, following the notation used by Hughes [7]. $N_a$ is used to denote the finite element shape function associated with node $a$. We use bilinear quadrilateral shape functions and compute all integrals using two-by-two Gauss quadrature.

The velocity equation (Eq 17) gives rise to the linear system

$$\mathbf{K}\mathbf{v}_{all} = \mathbf{F} \tag{S29}$$

The vector $\mathbf{v}_{all}$ contains the velocity values $[v_r, v_z]$ at each node, and the global stiffness matrix **K** and global load vector **F** are assembled from elemental contributions. The contributions from nodes $a$ and $b$ belonging to element $e$ are given as follows, where $\Omega_e$ denotes the element domain (note the factor of $2\pi r$ due to axial symmetry, as well as the term $(N_a N_b/r^2)$ which arises when computing $\nabla\mathbf{v}$ in cylindrical coordinates)

$$\begin{aligned}
\mathrm{K}_{arbr,e} &= \int\left(2\frac{\partial N_a}{\partial r}\frac{\partial N_b}{\partial r} + 2\frac{N_a N_b}{r^2} + \frac{\partial N_a}{\partial z}\frac{\partial N_b}{\partial z}\right)2\pi r d\Omega_e \\
\mathrm{K}_{arbz,e} &= \int\frac{\partial N_a}{\partial r}\frac{\partial N_b}{\partial z}2\pi r d\Omega_e \\
\mathrm{K}_{azbr,e} &= \int\frac{\partial N_a}{\partial z}\frac{\partial N_b}{\partial r}2\pi r d\Omega_e \\
\mathrm{K}_{arbr,e} &= \int\left(2\frac{\partial N_a}{\partial z}\frac{\partial N_b}{\partial z} + \frac{\partial N_a}{\partial r}\frac{\partial N_b}{\partial r}\right)2\pi r d\Omega_e
\end{aligned} \tag{S30}$$

For the load vector

$$\begin{aligned}
\mathrm{F}_{ar,e} &= \int p_{est}\left(\frac{\partial N_a}{\partial r} + \frac{N_a}{r}\right)2\pi r d\Omega_e + \int N_a \sigma_r 2\pi r d\Gamma_e \\
\mathrm{F}_{az,e} &= \int p_{est}\left(\frac{\partial N_a}{\partial z}\right)2\pi r d\Omega_e + \int N_a \sigma_z 2\pi r d\Gamma_e
\end{aligned} \tag{S31}$$

where the second term in the load vector arises from the Neumann boundary condition (stress balance) and $\Gamma_e$ denotes the boundary of the element. The vector $[\sigma_r, \sigma_z]$ denotes the boundary stresses given by adding the adhesion stress, protrusion stress, and cortical stress, which are only non-zero for those elements which belong to the free boundary of the cell. For elements in contact with the substrate, there is no contribution from the no-slip boundary condition because the enforced velocity is equal to zero.

The pressure equation (Eq 18) gives rise to the linear system

$$\mathbf{Q}\,\mathbf{p}_{all} = \mathbf{G} \tag{S32}$$

The vector $\mathbf{p}_{all}$ contains the pressure value $p$ at each node. The global stiffness matrix **Q** and global load vector **G** are assembled from elemental contributions given by

$$\mathrm{Q}_{ab,e} = \int \left[ N_a N_b - \epsilon \left( \frac{\partial N_a}{\partial r}\frac{\partial N_b}{\partial r} + \frac{\partial N_a}{\partial z}\frac{\partial N_b}{\partial z} \right) \right] 2\pi r d\Omega_e \tag{S33}$$

$$\mathrm{G}_{a,e} = \int N_a \left[ p_{est} - \frac{\partial v_r}{\partial r} - \frac{v_r}{r} - \frac{\partial v_z}{\partial z} \right] 2\pi r d\Omega_e \tag{S34}$$

The expression for $\mathrm{G}_a$ incorporates the assumption that $\nabla p \cdot \mathbf{n}$ is zero along the cell boundary, as dictated by the boundary condition, and $\nabla^2 p = 0$ everywhere, which is satisfied exactly for an incompressible fluid. Derivatives of $[v_r, v_z]$ are computed using bilinear quadrilateral shape functions.

This system of equations is solved iteratively, as described in Methods. Examples of the fluid velocities and pressures obtained by the Brownian zipper model (Fig S6A) and the protrusive zipper model (Fig 6B) are included for illustration.

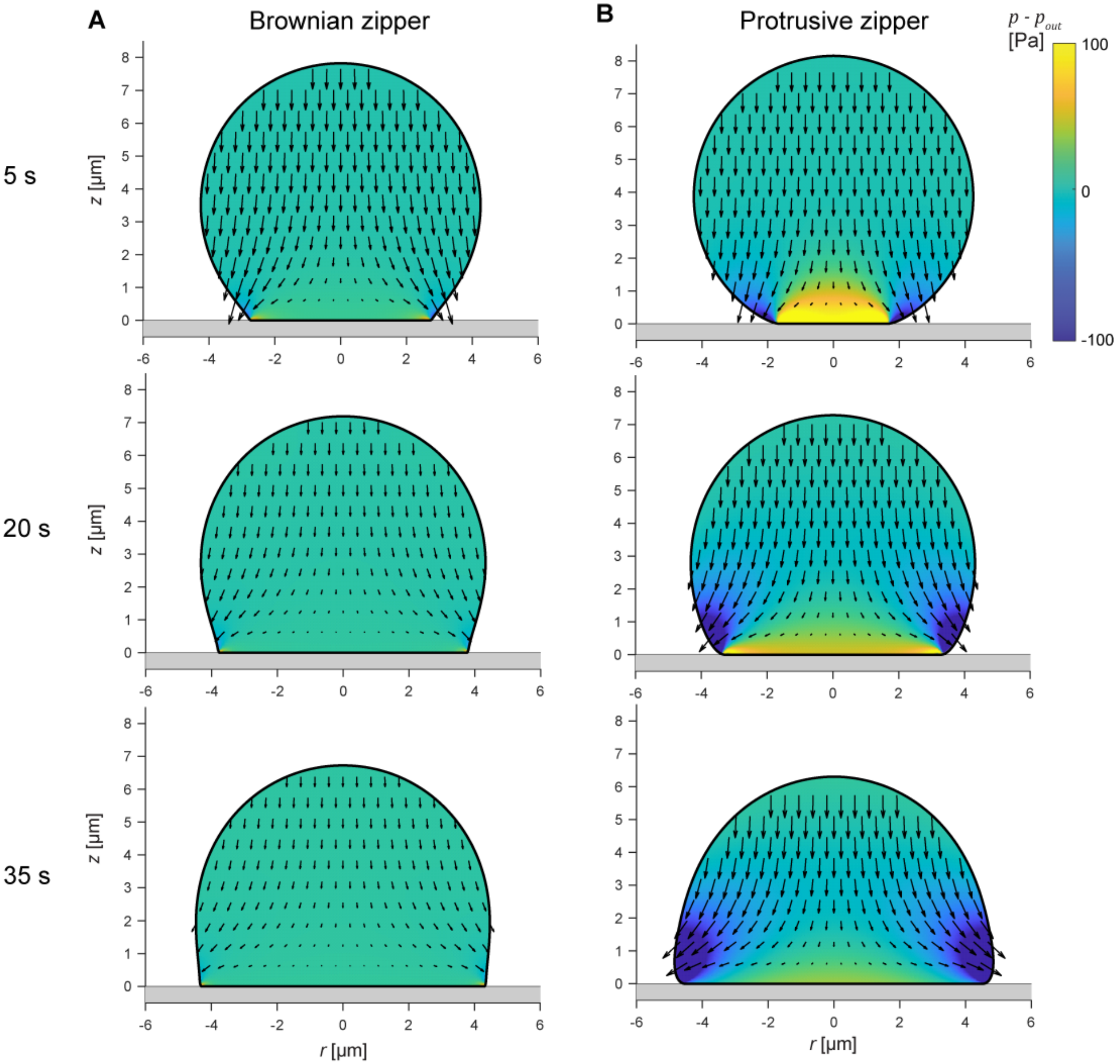


**Fig S6. Example velocity and pressure fields obtained by finite element computations.**
(A) Fluid velocity vector fields and pressure scalar fields (heat maps) are shown for purely adhesion-driven spreading with a ligand density of 10%. The magnitude of fluid velocity decays relatively quickly over time. (B) Fluid velocity vector fields and pressure scalar fields (heat maps) are shown for purely protrusion-driven spreading. The velocities maintain a similar magnitude over time due to the time-dependent increase in protrusion stress.

### Appendix E. Numerical testing and optimization

The accuracy of numerical computations such as used in this study can be affected by several factors. In this appendix, we document tests that we have carried out to verify the robustness of our numerical predictions. All tests of the Brownian zipper model are conducted for a ligand density ($\rho_l$) of 10%, and all tests of the protrusive zipper model are carried out in the absence of adhesion stress.

#### I. Test for incompressibility

Solving the governing equations for the perturbed Stokes equations given in Methods (Eqs 17 and 18) yields an approximate satisfaction of incompressibility. The degree to which incompressibility is enforced depends on the magnitude of the perturbation parameter $\epsilon$ in Eq 18. Specifically, as $\epsilon$ goes to zero, the fluid becomes more incompressible. However, if $\epsilon$ is too small, there are numerical difficulties. Thus, it is worthwhile to evaluate how closely incompressibility is satisfied for different values of $\epsilon$. In our model, $\epsilon$ scales with the characteristic radius of an individual element $h_{mesh}$ and the effective viscosity $\mu$ [8]. As described in Eq 15 of the main text, we generally require that $\epsilon \le h_{mesh}^2/\mu$ , but we can write more generally

$$\epsilon = \bar{\epsilon}\left(\frac{h_{mesh}^{2}}{\mu}\right) \tag{S35}$$

where $\bar{\epsilon}$ is a dimensionless scaling factor that should have a magnitude less than or equal to one to satisfy the above inequality. Here, we assess how this parameter affects the error in our model.

We quantify the degree of compressibility at different locations in the cell body in terms of the local rate of dilatation, obtained by integrating $\nabla \cdot \mathbf{v}$ over individual elements and dividing by the element volume $V_{el}$. That is,

$$\text{Dilatation rate} = \frac{\int(\nabla\cdot\mathbf{v})\,dV_{el}}{V_{el}} \tag{S36}$$

This dilatation rate has units of inverse seconds and describes the rate of fluid expansion per unit volume. In general, we expect the largest departures from perfect incompressibility to occur in the immediate vicinity of the perimeter of the contact region, due to a well-known singularity that has been examined thoroughly for different cases of droplet spreading [4]. Focusing on this region, we have characterized how our choice of the value of $\epsilon$ affects the dilatation rate (Fig. S7A). We evaluate the overall error of the model in terms of the mean square error (MSE) for the incompressibility condition

$$\text{MSE} = \frac{\int(\nabla\cdot\mathbf{v})^{2}\,dV}{V_{tot}} \tag{S37}$$

The MSE initially depends only weakly on the value of $\bar{\epsilon}$, but increases considerably at higher values, as expected (Fig S7B-C). Generally, a value of $\bar{\epsilon} = 1$ results in sufficiently low error while also avoiding numerical difficulties associated with lower values of $\bar{\epsilon}$.

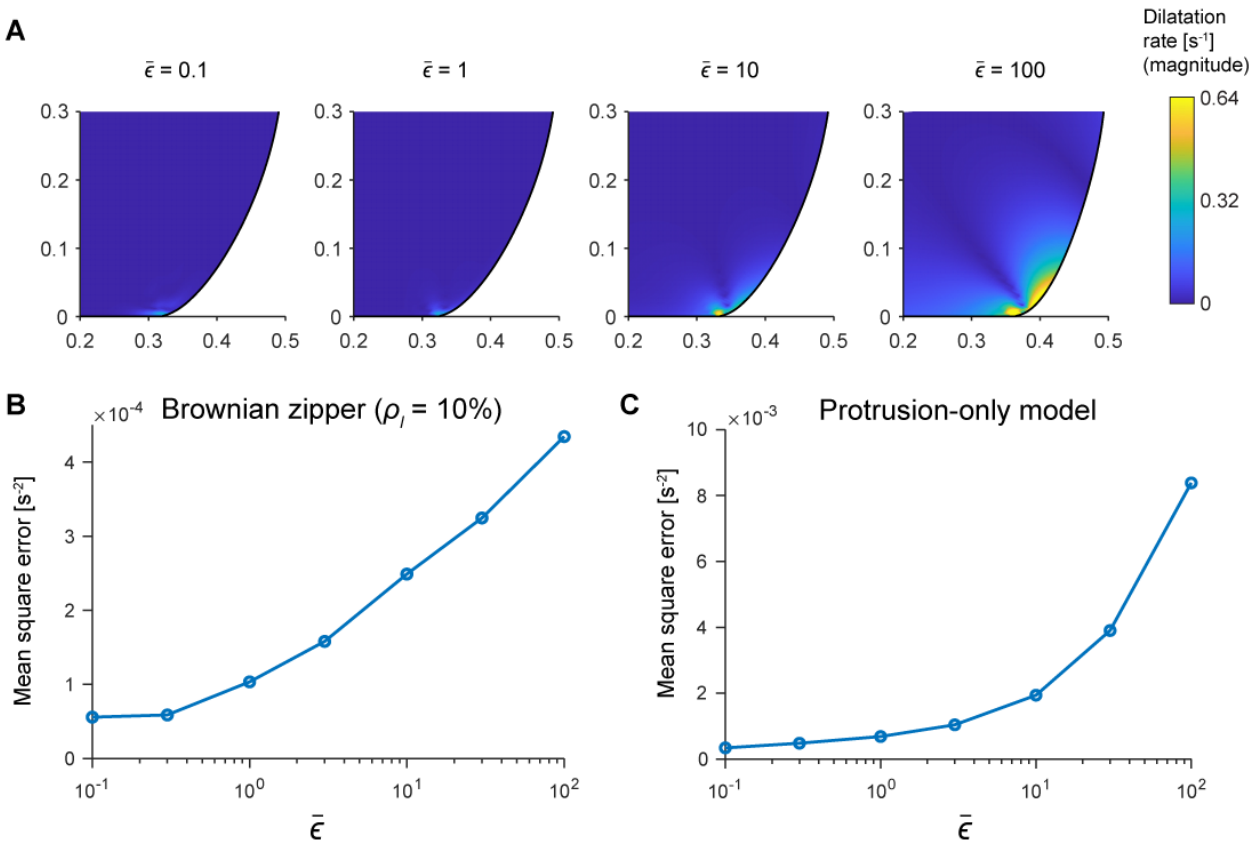


**Fig S7. Testing for incompressibility.**
(A) Heat maps of the dilatation rate within the leading edge of the cell for purely protrusion-driven spreading at $t$ = 15 s for different values of $\bar{\epsilon}$. (B) Volumetric mean square error is plotted as a function of $\bar{\epsilon}$ for Brownian Zipper simulations at $t$ = 15 s. (C) Volumetric mean square error is plotted as a function of $\bar{\epsilon}$ for protrusion-only spreading at $t$ = 15 s.

### II. Time-step testing

The optimal time step $\Delta t$ for fluid mechanics simulations depends on fluid velocity and the mesh resolution. A common reference for this is the Courant number, generally given by

$$C = \frac{v\Delta t}{l} \tag{S38}$$

where $l$ is a characteristic length and $v$ is a characteristic velocity. For numerical stability, $C$ should be much less than one.

We use Eq (S38) to determine the appropriate time step for the minimum ratio of $l/v$ across all elements; that is

$$\Delta t = C \min\left(\frac{l}{v}\right) = C \min_{\text{all el}}\left(\min\left[\frac{r_{span}}{v_{r,c}}, \frac{z_{span}}{v_{z,c}}\right]\right) \quad \text{(S39)}$$

where $r_{span}$ is the span of the element in the $r$ direction, $z_{span}$ is the span of the element in the $z$ direction, $[v_{r,c}, v_{z,c}]$ is the velocity vector at the centroid of the element, and $C$ is the desired Courant number. For instance, if $C = 0.1$, fluid flow crosses at most 10% of any individual element during a single time step. We performed a series of simulations using decreasing values of $C$ to establish at which Courant number our computations converge satisfactorily. Figure S8 shows examples of testing different time steps for the Brownian Zipper model (Fig S8A) and the protrusive zipper model with no adhesion stress (Fig S8B). For smaller time steps, the contact area curves become smoother and converge to a single solution. Generally, we find that $C \le 0.1$ performs well in these cases, which is the value we choose for our simulations.

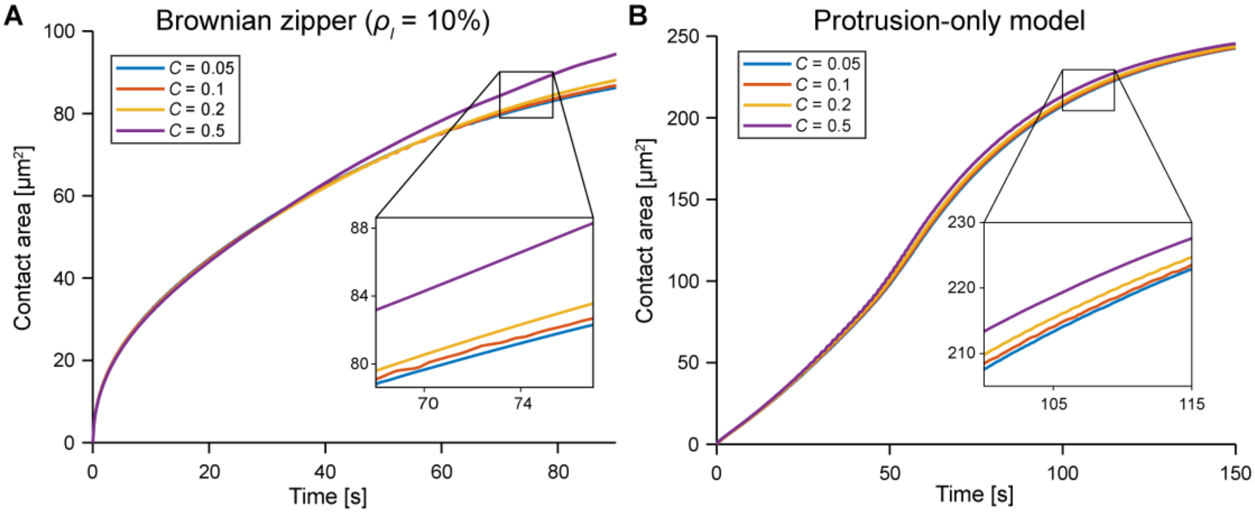


**Fig S8. Time-step testing.**
Contact area versus time for different Courant numbers for (A) the Brownian zipper model and (B) purely protrusion-driven spreading.

### III. Mesh refinement

We ensured that the results of our simulations do not depend significantly on the mesh resolution as follows. When constructing the mesh, we start by specifying the boundary nodes, choosing tight spacing near the substrate and gradually increasing spacing to 0.17 μm over the rest of the cell contour (Fig S9A). After parameterizing the contour, we use transfinite interpolation to construct the interior mesh ([9], Fig S9A). Using a global edge spacing of 0.17 μm generally results in about 2,000 elements. Here, we focus on refining the mesh spacing close to the surface where the stresses are most concentrated.

For both the Brownian zipper model and the protrusive zipper model, we tested minimum boundary point spacing values $\Delta s_{min}$ of 25 nm, 10 nm, 5 nm, and 2.5 nm. In both cases, we find convergence for $\Delta s_{min} \leq 5$ nm (Fig S9B-C). For our simulations in the main paper, we use $\Delta s_{min}$ = 2.5 nm.

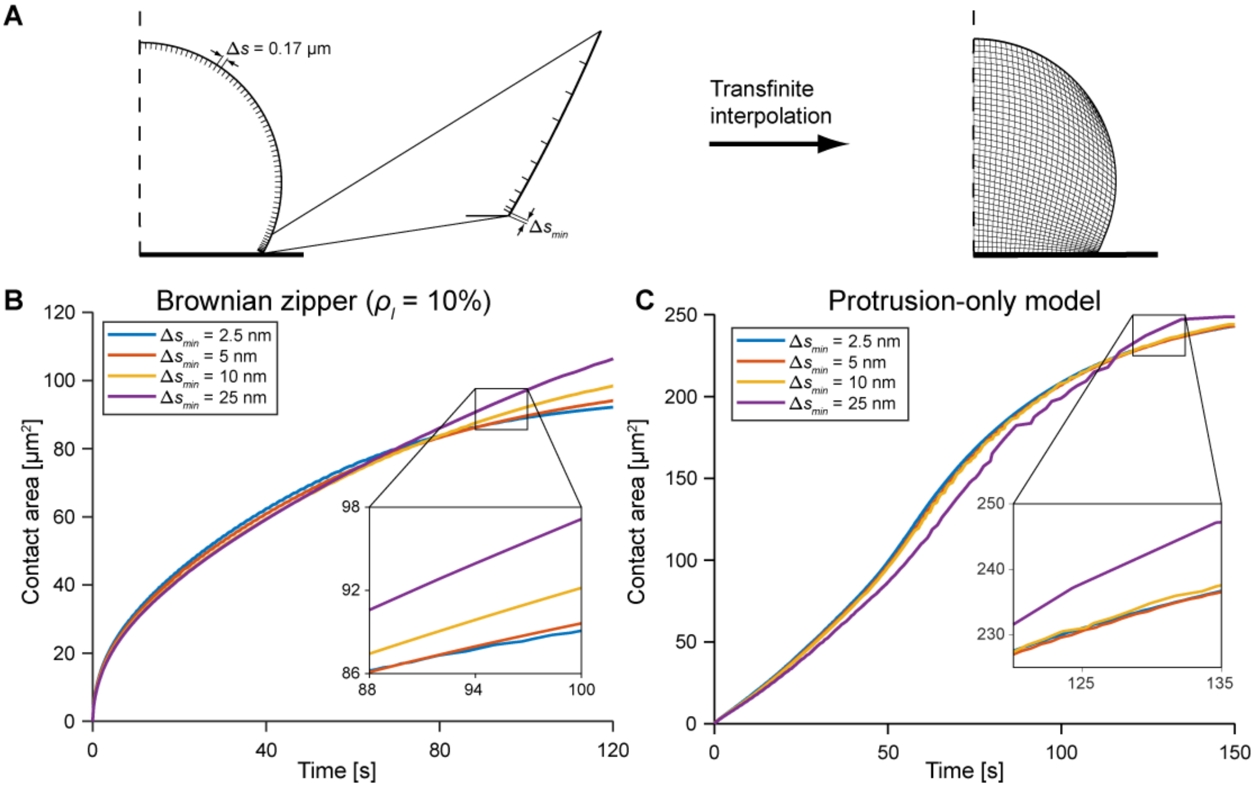


**Fig S7. Mesh refinement.**
(A) Illustration of mesh construction including our choice of relevant boundary spacing intervals. Convergence of the simulations is evaluated by inspection of contact-area-versus-time curves obtained with different minimum mesh spacings close to the substrate ($\Delta s_{min}$) for (B) the Brownian zipper model and (C) purely protrusion-driven spreading.

### Appendix F. Additional testing of the discrete adhesion model

As outlined in the main text, we introduced two main rules to implement a version of our protrusive zipper with discrete adhesion sites: 1) If the cell membrane is less than a threshold distance from a ligand (set by the magnitude of membrane fluctuations dictated by Eq 12), the membrane is considered bound, and 2) the protrusion stress decays as a function of time since last binding a ligand (Eq 13). The second rule contains a free parameter $t_0$, corresponding to a characteristic decay time for the protrusion stress. Because this parameter is not directly quantifiable from our data, we sought to determine how sensitive the outcomes of this version of the model were to our choice of $t_0$ (Fig S10).

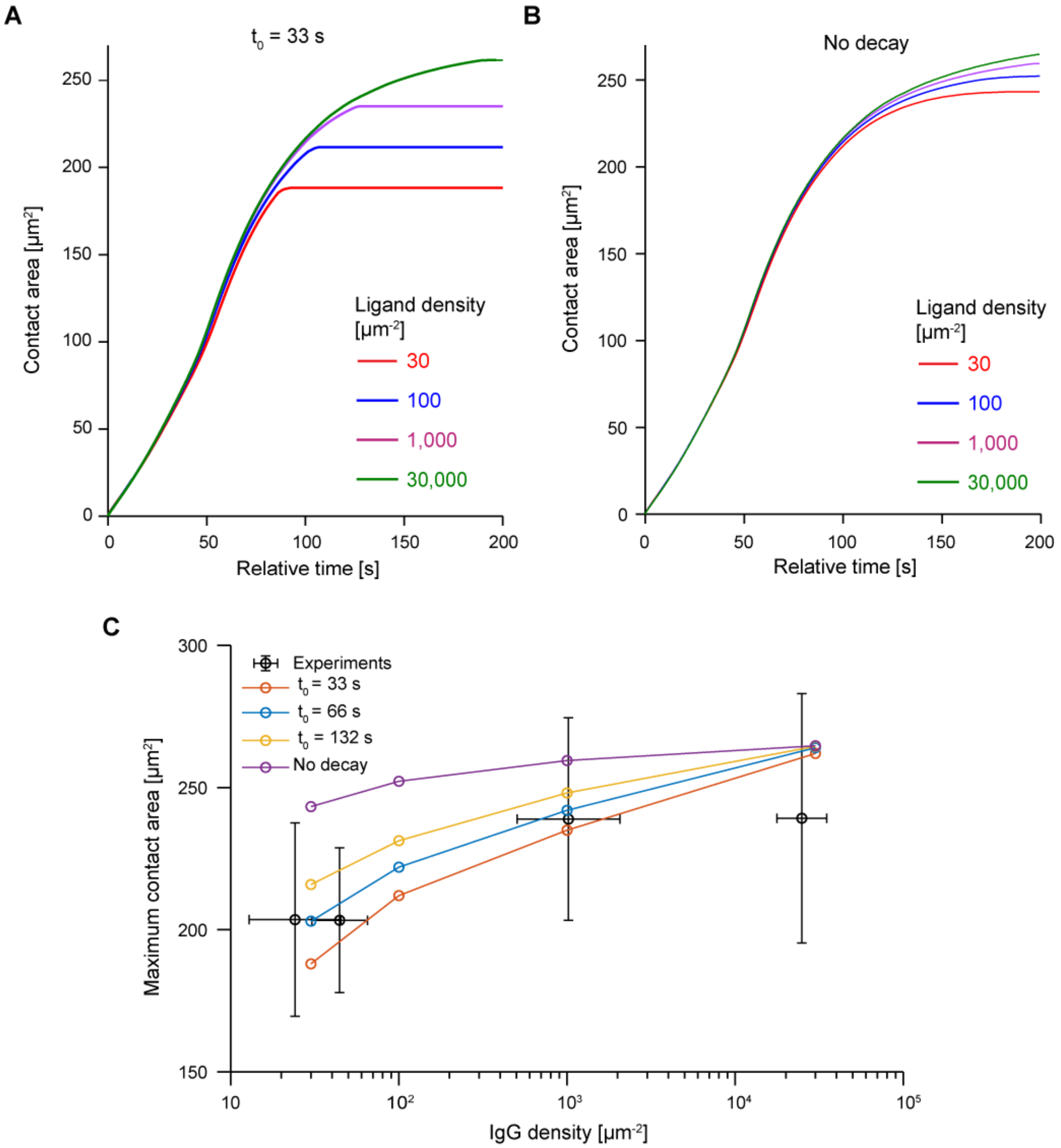


**Fig S10. Effect of varying $t_0$ on the protrusive zipper with discrete adhesion sites.**
Contact-area-versus-time for spreading over different densities of discrete binding sites, plotted for $t_0$ = 33 s (faster decay) (A) and for no time-dependent decay of the protrusion stress (B). (C) The relationship between maximum contact area and IgG density depends on how quickly the protrusion stress decays.

In the main text, we use $t_0 = 66$ s, and here we tested $t_0$ with either half this value (33 s, faster decay) or twice this value (132 s, slower decay). We also tested the model without any decay by setting $t_0 = \infty$. Regardless of the choice of $t_0$, we observe the same qualitative behavior of the model – spreading speeds remain similar across different ligand densities, but maximum contact area increases as a function of ligand density (Fig S10A,B). However, the exact relationship between ligand density and maximum contact area depends on the choice of $t_0$. Faster decay rates (lower $t_0$) generally resulted in earlier termination of spreading, but with slow or absent decay, cells continue spreading longer (Fig S10). This difference is most evident at lower ligand densities, where a more sustained protrusion stress is required to overcome larger gaps between ligands.

We also noted that our experiments seemed to indicate a saturation in maximum contact area for IgG densities greater than about 1,000 $\mu m^{-2}$, but our model predicts a continued increase in maximum contact area up to the highest density tested (30,000 $\mu m^{-2}$). At these higher densities of IgG, receptors are the limiting factor of the maximum adhesion strength, which is not accounted for in our model. Therefore, we conducted additional proof-of-principle simulations in which both ligands and receptors were treated as discrete.

On the order of $1 \times 10^6$ Fcγ receptors are present in the membrane of a passive neutrophil [10, 11]. The total membrane surface area includes not just the apparent surface area, but also membrane folds such as microvilli. Because neutrophils can expand their apparent surface area up to three times its resting value [12, 13], we conservatively estimated the total receptor density $\rho_{Fc\gamma R}$ as $1 \times 10^6 / 3SA_0$, or about 1,470 receptors per square micron.

Because these receptors are generally mobile, we assumed that any given receptor effectively occupied a length of membrane $\delta l$ dictated by receptor mobility (effective diffusion coefficient $D_{eff}$) and the current time step $\Delta t$:

$$\delta l = \sqrt{4 D_{eff} \Delta t} \qquad \text{(S40)}$$

If a given receptor is centered at a point on the membrane specified by arc length $s_R$, then the receptor can diffuse within a range bounded by $s_R - \delta l$ and $s_R + \delta l$ during a time interval of $\Delta t$ (Fig S11A inset). We here postulate that, for any given ligand, if the minimum distance to this receptor, $d_{min}$, is less than $d_{thresh}$ (given by Eq 12 in the main text), then binding occurs.

Using the above approach to discretize receptors, we require the total region over which each receptor can diffuse ($2\delta l$ from Eq (S40)) to be less than the receptor spacing; otherwise, the entire membrane would be available to bind free ligand, as in the original discrete adhesion model. This places an upper limit of about $4 \times 10^{-4}$ $\mu m^2$/s on $D_{eff}$, much lower than the values reported for Fcγ receptor diffusion coefficients (about 0.01 to 0.1 $\mu m^2$/s) [14]. However, $D_{eff}$ should be viewed as an effective parameter, capturing not only diffusion but also the effects of lateral confinement [15] and close contacts [16].

As a proof of concept, we performed simulations with $\rho_{Fc\gamma R} = 1{,}470$ $\mu m^{-2}$ and $D_{eff} = 1 \times 10^{-4}$ $\mu m^2$/s. (Fig S11) These simulations indeed showed that spreading on lower densities of IgG was similar to our original simulations, but spreading on the highest density terminated earlier due to limited receptor availability (Fig S11A,B). The results of this version of the model yielded an even better match to the experimentally measured relationship between IgG density and maximum contact area (Fig 8C).

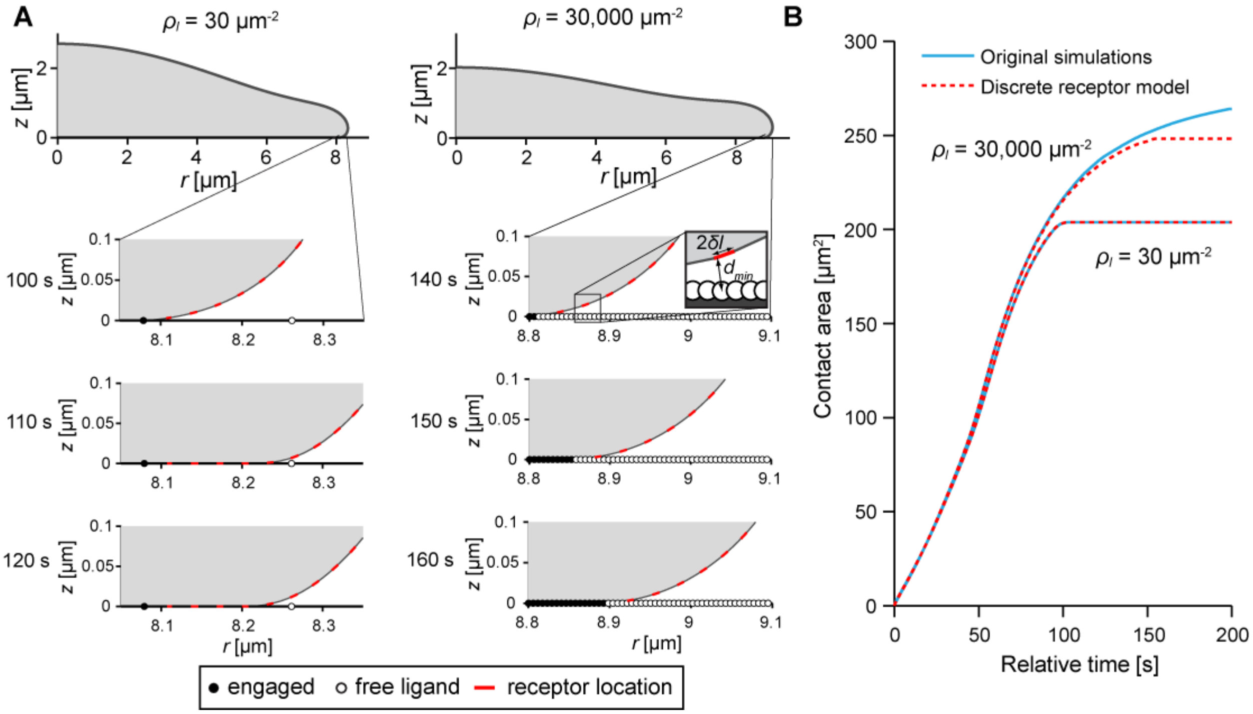


**Fig S11. Protrusive zipper with discrete ligands and discrete receptors.**
(A) Sample cell profiles for spreading over different densities of binding sites with receptors only present in the membrane segments shown in red. The inset defines the variables used in the text. (B) Spreading over low densities of IgG (30 µm$^{-2}$) is unaffected by a discrete receptor model, whereas spreading is receptor-limited on high densities of IgG (30,000 µm$^{-2}$).
