## Supplementary material for "Integrative experimental/computational approach establishes active cellular protrusion as the primary driving force of phagocytic spreading by immune cells": S7 Table (parameter values)

### S7 Table. Modeling parameters

| Parameter | Description | Value | Notes |
|---|---|---|---|
| $\tau_0$ | Resting cortical tension | 0.01 mN/m | Matches experimental value used by Herant et al [1] |
| $\tau_{max}$ | Maximum cortical tension | ~1 mN/m (see constitutive relation) | In a reasonable range for high tensions measured during neutrophil phagocytosis [2, 3] |
| $\mu$ | Effective cytoplasmic viscosity | 200 Pa-s (passive)<br>1,660 Pa-s (active) | Matches experimentally measured values [4, 5] |
| $\kappa_b$ | Effective membrane bending modulus | $1\times10^{-18}$ J | Matches experimentally measured values [6], much higher than values for RBCs or vesicles due to the neutrophil cortex |
| $R_0$ | Initial cell radius | 4.25 μm | Radius of a human neutrophil |
| $\sigma_0$ | Adhesion stress constant | 370 Pa | Together with the ligand density, this determines adhesion strength |
| $\rho_{IgG,max}$ | Ligand density corresponding to 100% for Brownian Zipper model | 10,000 IgG/$\mu m^2$ | This density corresponds to about 600 $\mu J/m^2$, which exceeds values derived from other cases of cell spreading [7] |
| $D_0$ | Zero adhesion force distance | 50 nm | Relatively large distance required for mesoscopic model, as standard in other continuum models [8, 9] |
| $\sigma_{prot,max}$ | Max. protrusion stress | 3,500 Pa | Actin filaments growing in parallel can achieve forces above 1 nN per $\mu m^2$ (> 1 kPa) [10, 11] |
| $s_0$ | Protrusive force range | 0.8 μm | Controls how the protrusion stress decays along the membrane (Eq 8) |
| $t_0$ | Characteristic time for decay of protrusion stress | 66 s | Used for discrete adhesion model (Eq 13) |
| $k_BT$ | Energy scale factor | $4.11\times10^{-21}$ J | Boltzmann constant ($k_B$) times room temperature (298 K), sets scale for membrane fluctuations (Eq 12) and ligand-receptor binding energy. |
| $\rho_{Fc\gamma R}$ | FcγR density in the neutrophil membrane | 1,470 $\mu m^{-2}$ | Receptor density used in discrete ligand + discrete receptor simulations shown in Fig 8B, explained in Appendix F |
| $D_{eff}$ | Effective FcγR diffusion coefficient | $1\times10^{-4}$ $\mu m^2/s$ | Chosen value explained in Appendix F |
